# Soluble cyclase-mediated nuclear cAMP synthesis is sufficient for cell proliferation

**DOI:** 10.1101/2022.05.18.492464

**Authors:** Alejandro Pizzoni, Xuefeng Zhang, Nyla Naim, Daniel L Altschuler

## Abstract

cAMP is a key player in many physiological processes. Classically considered to originate solely from the plasma membrane, this view was recently challenged by observations showing that GPCRs can sustain cAMP signaling from intracellular compartments associated with nuclear PKA translocation and activation of transcriptional events. In this report we show that neither PKA translocation nor cAMP diffusion, but rather nuclear sAC activation represents the only source of nuclear cAMP accumulation, PKA activation, and CREB phosphorylation. Both pharmacological and genetic sAC inhibition, that did not affect the cytosolic cAMP levels, completed blunted nuclear cAMP accumulation, PKA activation and proliferation, while an increase in sAC nuclear expression significantly enhanced cell proliferation. Moreover, utilizing novel compartment-specific optogenetic actuators we showed that light-dependent nuclear cAMP synthesis can stimulate PKA, CREB and trigger cell proliferation. Thus, our results show that sAC-mediated nuclear accumulation is not only necessary but sufficient and rate-limiting for cAMP-dependent proliferation.

## INTRODUCTION

Cyclic adenosine monophosphate (cAMP), the first second messenger described ^1,2^, is a key intermediate in signaling pathways controlling many cellular processes including proliferation, differentiation, survival, and metabolism ^3^. The players involved in its synthesis (adenylyl cyclases, ACs) and degradation (phosphodiesterases, PDEs) are relatively well-defined; however, the mechanisms cells utilize to compute (code/decode) the relay of the cAMP signal (*i.e*., fidelity, specificity, efficiency) are still not understood.

In the canonical cAMP pathway, an activated GPCR (G-protein coupled receptor) couples to heterotrimeric Gs to stimulate a transmembrane adenylyl cyclase (tmAC, AC1-9) at the plasma membrane; synthesized cAMP directly binds and activates a set of effectors ^4–10^, including protein kinase A (PKA), whose catalytic subunit (PKA-C) can diffuse to the nucleus, phosphorylate transcription factors and initiate the transcription of cAMP-specific genes ^11^,^12^. An additional intracellular source of cAMP is soluble adenylyl cyclase (sAC, AC10) which, unlike tmACs that are regulated by Gs and forskolin (FK), is activated by bicarbonate, Ca^2+^, and ATP ^13^. In addition to the cytosol, sAC was found at discrete cellular locations including the nucleus ^14^, where it represents the only cAMP-synthesizing activity. These findings showed that cAMP can be synthesized deep in the cell and ruled out the plasma membrane as the only cAMP source.

TSHR (Thyroid-stimulating hormone receptor) is a class A GPCR activated by TSH, a glycoprotein synthesized by the thyrotrophs in the anterior lobe of the pituitary (adenohypophysis) that is the main regulator of the thyroid gland ^15^. TSHR activation leads to membrane G-protein activation and stimulation of secondary messenger pathways, mainly Gs-AC-cAMP ^16,17^. Downstream activation of cAMP effector pathways, eg Epac1 and PKA, work synergistically towards thyrocyte proliferation ^18,19^, establishing thyroid/TSHR as a bona fide model to study cAMP signaling.

The spatiotemporal properties of signaling intermediates impart specificity on biological outputs ^20–24^. According to the classical view, cAMP signaling was considered to originate solely from the plasma membrane, with receptor endocytosis mediating signal termination, leading to a transient cAMP response ^25^. This original view was recently challenged by observations, first reported for TSHR ^26^, showing that upon internalization GPCRs can sustain cAMP signaling from intracellular membranes upon internalization. In this novel “two-wave paradigm”, upon the initial transient plasma membrane-generated cAMP wave, a sustained second wave is observed following agonist-induced receptor internalization into the endocytic compartment ^27–33^. This temporal bias might convey informational content if cells are able to discriminate cAMP production from different sources and trigger unique biological outcomes differentially associated with these two phases. In this context, a sustained cAMP elevation was reported to be specifically required in several physiological responses, such as chronic inflammation and pain ^34–37^, to resume meiosis in the oocyte ^38^, and several reports indicate its role in cAMP-dependent nuclear transcription ^30,31,39–43^. Similarly, internalization-mediated sustained cAMP elevation can aid in producing stable responses to pulsatile or low-concentration hormones such as PTH. In this respect, fast cytosolic and nuclear cAMP elevations were observed upon the synergistic action of adrenergic and PTHR receptors in an internalization-dependent manner ^39^. More conclusively, it was recently demonstrated that only a photoactivated bacterial cyclase (bPAC) targeted to the outer mitochondria or endosome but not to the plasma membrane can induce a light-dependent CREB phosphorylation and nuclear transcription ^42^. Thus, it is becoming clear that endocytosis, via altering the subcellular location of cAMP production rather than changing the total amount of cAMP produced, might facilitate the selective activation of distinct PKA pools ^40^,^43^. Furthermore, recent studies suggest the existence of a pool of PKA holoenzyme in the nucleus ^39,44,45^, adding another challenge to the classical cAMP model.

The existence of a buffering capacity ^46,47^ and of PDE nanodomains ^46^ at different cellular locations can explain the ‘barriers’ that prevent cAMP produced at different locations from transducing its full repertoire of downstream effects; for example, it is known that in some cells plasma membrane generated cAMP is unable to reach the nuclear compartment ^48^, unless PDE activity is inhibited ^42^. Moreover, the kinetic discrepancies between nuclear cAMP accumulation and nuclear PKA activation also indicated the existence of cAMP nanodomains where AKAP-PKA-PDE complexes constrain cAMP levels and control nuclear PKA activation ^44^. Thus, despite the controversy of whether cAMP diffusion or PKA-C translocation represents the ‘rate-limiting step’, it is becoming increasingly clear that the location, localization, and duration of signals are key factors determining nuclear PKA activation and cAMP-dependent transcription. However, whether cAMP diffuses from the cytosol or is synthesized locally in the nucleus is still unknown.

In the present study, we demonstrate using a combination of pharmacological, genetic, and optogenetic manipulations that local sAC activation is the source of TSH-mediated nuclear cAMP production/elevation. While sAC inhibition minimally affected the cytosolic cAMP levels, it completed blunted nuclear cAMP accumulation and PKA activation. Moreover, our results show that sAC-mediated nuclear accumulation is not only necessary but sufficient and rate-limiting for cAMP-dependent proliferation.

## RESULTS

### TSH increases nuclear cAMP in an sAC-dependent manner

To monitor cAMP dynamics in live cells, we exploited the TSH/PCCL3 thyroid model system we have extensively used in the past ^18,19,49–52^. Real-time cAMP recordings were performed by transfecting PCCL3 thyroid follicular cells with cytosolic and nuclear-targeted FRET-based cAMP sensors, as we reported before ^50^ (Fig. S1a). As shown in Figs. 1a and 1b, TSH stimulated a sustained cAMP increase in both cytosolic and nuclear compartments. Nuclear events are frequently associated with a sustained elevation of cytosolic cAMP that is dependent on dynamin-mediated endocytosis. Accordingly, pharmacological (Dyngo 4a and Dynasore) and genetic (dn K44A) dynamin inhibitors, blocked the TSH-mediated sustained phase in a dose-dependent manner and blunted nuclear cAMP accumulation (Fig.1c-e and S1b-f).

**Fig. 1:**
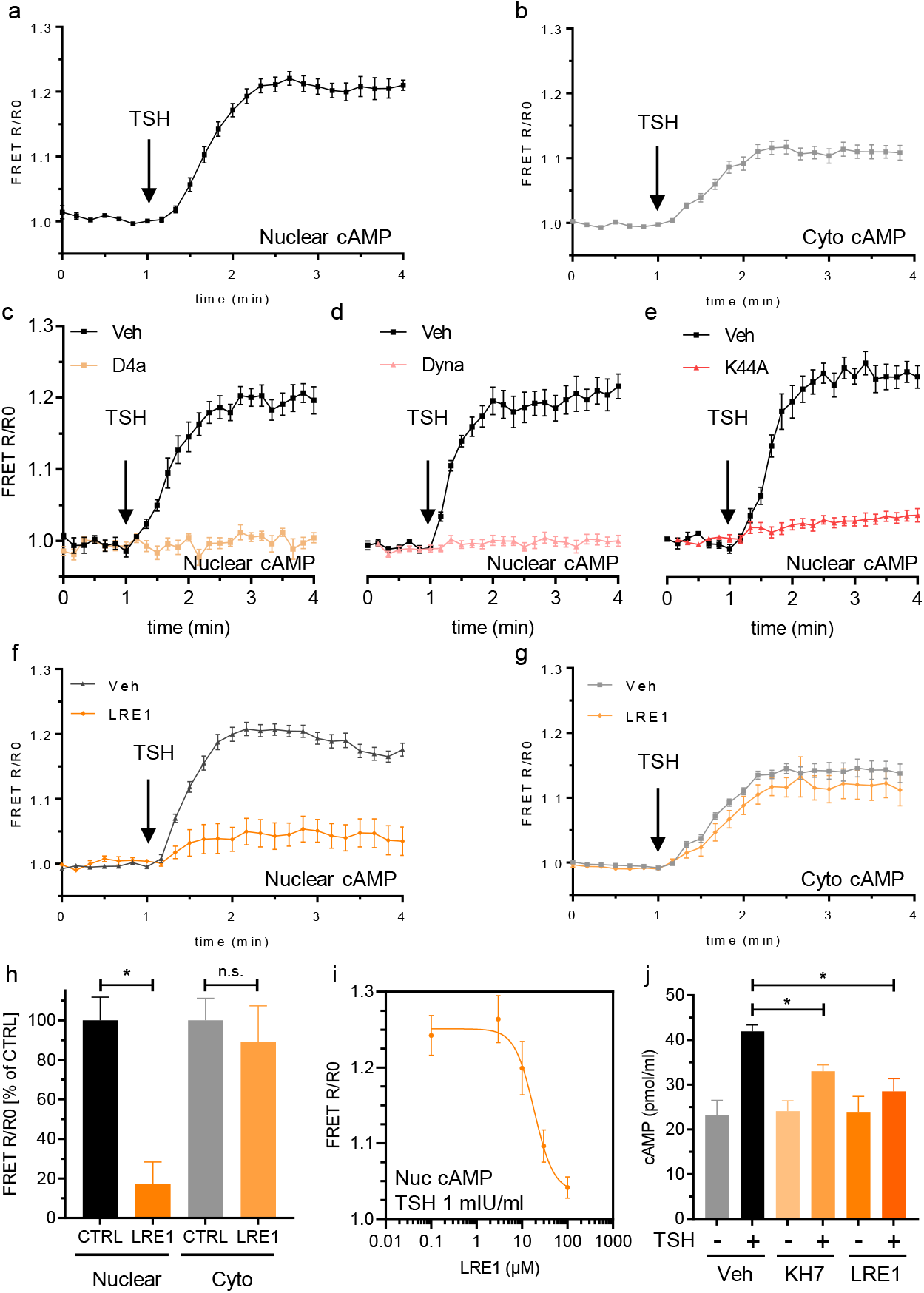
TSH elevates nuclear cAMP in an sAC-dependent manner. Real-time FRET-based monitoring of cAMP levels in live PCCL3 cells stimulated with TSH (1 mIU/ml). Traces are the normalized FRET ratios (R/R0) of the NLS-H208 (nuclear; a) and H208 (cytosolic; b) sensors. Nuclear cAMP measurements of cells preincubated for 5 minutes with Dyngo 4a (D4a, 30 μM; c) or Dynasore (Dyna, 30 μM; d). Nuclear cAMP measurements with or NLS-H188 (FRET sensor) in cells transfected with the mutant Dynamin K44A (K44A, e). Nuclear (f) and cytosolic (g) AMP measurements after 30 min of incubation with an sAC inhibitor (LRE1; 100 μM) or Vehicle (Veh; DMSO). Normalized ratios are represented as the % of its respective controls (FRET R/R0 [% of CTRL]) after 5 minutes of incubation with TSH (h). Concentration-dependent inhibition of the TSH response by LRE1. (i). Cyclase activity and cAMP steady-state measurements (pmol/ml) were performed after labeling the cellular pool of ATP with [^3^H]-adenine. Separation of [^3^H]-cAMP product by sequential chromatography in Dowex/Alumina was performed in TSH-stimulated (+) cells or unstimulated (-) cells pretreated either with vehicle (Veh; DMSO), KH7 (100 μM), or LRE1 (100 μM) (j). Data are expressed as mean ± SEM (for n cells/samples and independent experiments, see statistics). Significance was tested using a two-tailed Student’s t-test (*, p<0.05; n.s., not significant).

The increase in nuclear cAMP could represent diffusion from the cytosol or local synthesis by nuclear sAC. sAC is expressed in PCCL3 cells, as evaluated by immunoblotting and immunofluorescence, confirming its presence in the nucleus (Fig. S2a-b). To evaluate the role of sAC in TSH-mediated nuclear cAMP accumulation, we preincubated cells with LRE1, a selective sAC inhibitor that does not affect the activity of tmACs ^53^. While LRE1 minimally affected the cytosolic component (< 10%), it significantly inhibited nuclear cAMP accumulation (~ 80%) (Figs. 1f-h, and S2c). Dose-responses showed that LRE1 inhibits nuclear cAMP signals with an EC50 ~ 18μM (Figs. 1i and S1d). To validate the FRET results, total cAMP was assessed biochemically from cell lysates preincubated with LRE1 and KH7, another sAC selective inhibitor ^54^, confirming a component of sAC activity in TSH-mediated total cAMP accumulation (Fig. 1h). Similar results were obtained for β2-adrenergic receptor (β2-AR) in HC-1 cells. HC-1 cells endogenously express β2-AR and Gs, but they do not respond to isoproterenol ^55^ unless transfected with a tmAC ^56^. sAC expression in untransfected HC-1 cells was confirmed by the LRE1 sensitivity of the bicarbonate-mediated nuclear cAMP accumulation (Fig. S3a-d). Transfection of AC9, rescued the cytosolic isoproterenol response, accompanied by a nuclear cAMP accumulation that was sensitive to LRE1 and Dynasore (Fig. S4a-c). Thus, these combined results indicate that stimulation of both β2-AR (in HC1 cells) and TSHR (in PCCL3 cells), trigger a nuclear sAC-mediated cAMP accumulation that is dependent on receptor internalization.

### sAC activity is required and rate-limiting for cell proliferation

Next, we decided to test whether sAC inhibition has an impact on TSH-mediated cell proliferation. Pharmacological inhibition of sAC by LRE1 or KH7 blocked cell proliferation, assessed by both ^3^H-thymidine and BrdU incorporation assays (Fig. 2a). To confirm the pharmacological results, we generated an shRNA for rat sAC. sh-sAC-mediated sAC down-regulation but not sh-vector, blocked TSH-mediated cell proliferation (Fig. 2b). sAC 1-469 (sACt) is a truncated and highly active soluble cyclase variant ^57^. Mutations were introduced to generate a sh-resistant sACt (sACt^R^) and immunofluorescence staining confirmed both sACt^R^ and sACt are present in cytosol and nucleus (Figs. S5a-b); however, only expression of sACt^R^ but not sACt was able to rescue sh-sAC-mediated inhibition (Fig. 2b). Interestingly, expression of sACt^R^/sACt consistently showed a proliferative response in TSH-stimulated samples (~ 60% BrdU/tag) above the usual values observed in non-transfected or sh-vector controls (~ 30-40% BrdU/tag). This observation prompted us to study this effect in more detail confirming that expression of sACt^R^/sACt increased TSH-mediated cell proliferation in a dose-dependent manner without losing the sensitivity to LRE1 (Fig. 2c). Moreover, consistent with the pharmacological inhibition by LRE1, sAC knockdown, and sACt overexpression affected TSH-mediated nuclear cAMP accumulation without interfering with the cytosolic cAMP response (Figs. 2d-e, and S5c). Thus, these results confirmed that sAC activity is required for the TSH-mediated proliferative response and suggest that local sAC-generated cAMP signals in the nucleus could represent a rate-limiting step for cell proliferation.

**Fig. 2:**
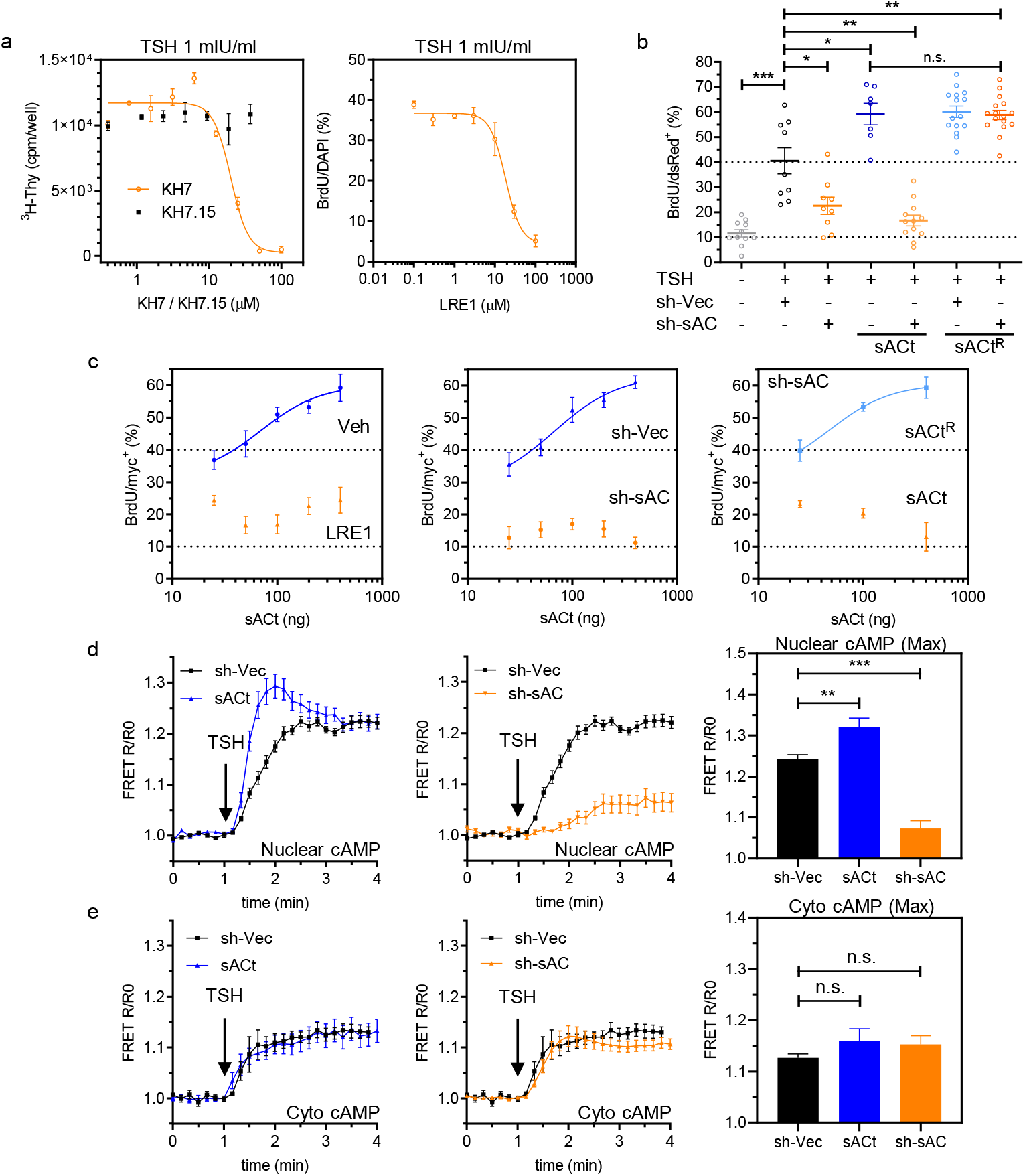
sAC activation is necessary and rate-limiting for G1/S progression. (a) Concentration-dependent inhibition of the TSH-triggered (1 mIU/ml) proliferation in PCCL3 cells by KH7, KH7.15, and LRE1. Proliferation was assessed by ^3^H-thymidine (left panel) and BrdU (right panel) incorporation. PCCL3 cells were incubated for 30 min with inhibitors before stimulation with TSH. ^3^H-thymidine data are expressed as the averaged counts per minute measured in each well (cpm/well). (b) Cells were transfected with pSiren-red-sh-sAC1 (sh-sAC) or empty vector (sh-Vec) and BrdU labeling was used to monitor TSH-dependent G1/S progression in the presence of 5% FBS as co-mitogen. The inhibitory response can be rescued with a sh-resistant sAC (sAC^R^) but not with sACt. BrdU data are expressed as % BrdU/ dsRed^+^). Broken lines represent the average %BrdU/DAPI values of control and TSH-stimulated parental cells. (c) sACt expression increases proliferation. Cells were transfected with increasing doses (25, 50, 100, 200, and 400 ng) of sACt/sACt^R^ plasmids, and proliferation was assessed using BrdU incorporation. Experiments were performed in cells transfected sACt in the presence of vehicle (Veh; DMSO) or LRE1 (100 μM) (left panel) and in cells transfected with sACt and either sh-Vec or sh-sAC (middle panel). Cells were also transfected with sh-sAC and either sACt or sACt^R^ (right panel). (d) Real-time FRET-based monitoring of nuclear cAMP levels in live PCCL3 cells. Traces are the normalized FRET ratios (R/R0) of the NLS-H208 (nuclear) cAMP sensor. Nuclear cAMP measurements were performed in cells transfected with sACt or sh-Vec (left panel) and with sh-sAC or sh-Vec (middle panel). Quantitative analysis of maximum (Max) nuclear R/R0 values during the initial 5 min (right panel). (e) Cytosolic cAMP measurements (H208) were performed in cells transfected with sACt or sh-Vec (left panel) and with sh-sAC or sh-Vec (middle panel). Quantitative analysis of maximum (Max) cytosolic R/R0 values during the initial 5 min (right panel). Data are expressed as mean ± SEM (for n cells/samples and independent experiments, see statistics). Significance was tested using a two-tailed Student’s t-test Student’s t-test (*, p<0.05; **, p<0.01; ***, p<0.001; n.s., not significant).

### Nuclear sAC activity is the rate-limiting step for cell proliferation

In order to assess whether sAC expression in the nuclear compartment is relevant for cell proliferation, we transfected PCCL3 cells with either nuclear (NLS) or plasma membrane (Lyn) targeted versions of a mCherry-tagged sACt ^45^. Like the non-targeted version of sACt (Fig. 2b), NLS-sACt but not Lyn-sACt showed a dose-dependent effect on TSH-mediated proliferation (Figs. 3a and S6a). Like LRE1 and sh-sAC, CRISPR/Cas9-mediated sAC deletion (Figs. S6b-d) blocked TSH-mediated proliferation and reduced nuclear but not cytosolic cAMP dynamics (Figs. 3b and 3c). Only a sg-resistant NLS-sACt (NLS-sACt^R^) was able to rescue sg-sAC-mediated inhibition, and like untagged sACt^R^/sACt (Fig. 2b), NLS-sACt^R^/NLS-sACt expression consistently showed a high proliferative response in TSH-stimulated samples (Fig. 3b). Thus, these results show that a nuclear-targeted sAC can sustain cell proliferation even in an sAC-knockdown context and indicate that nuclear sAC activation is a limiting factor controlling TSH-mediated proliferation.

**Fig. 3:**
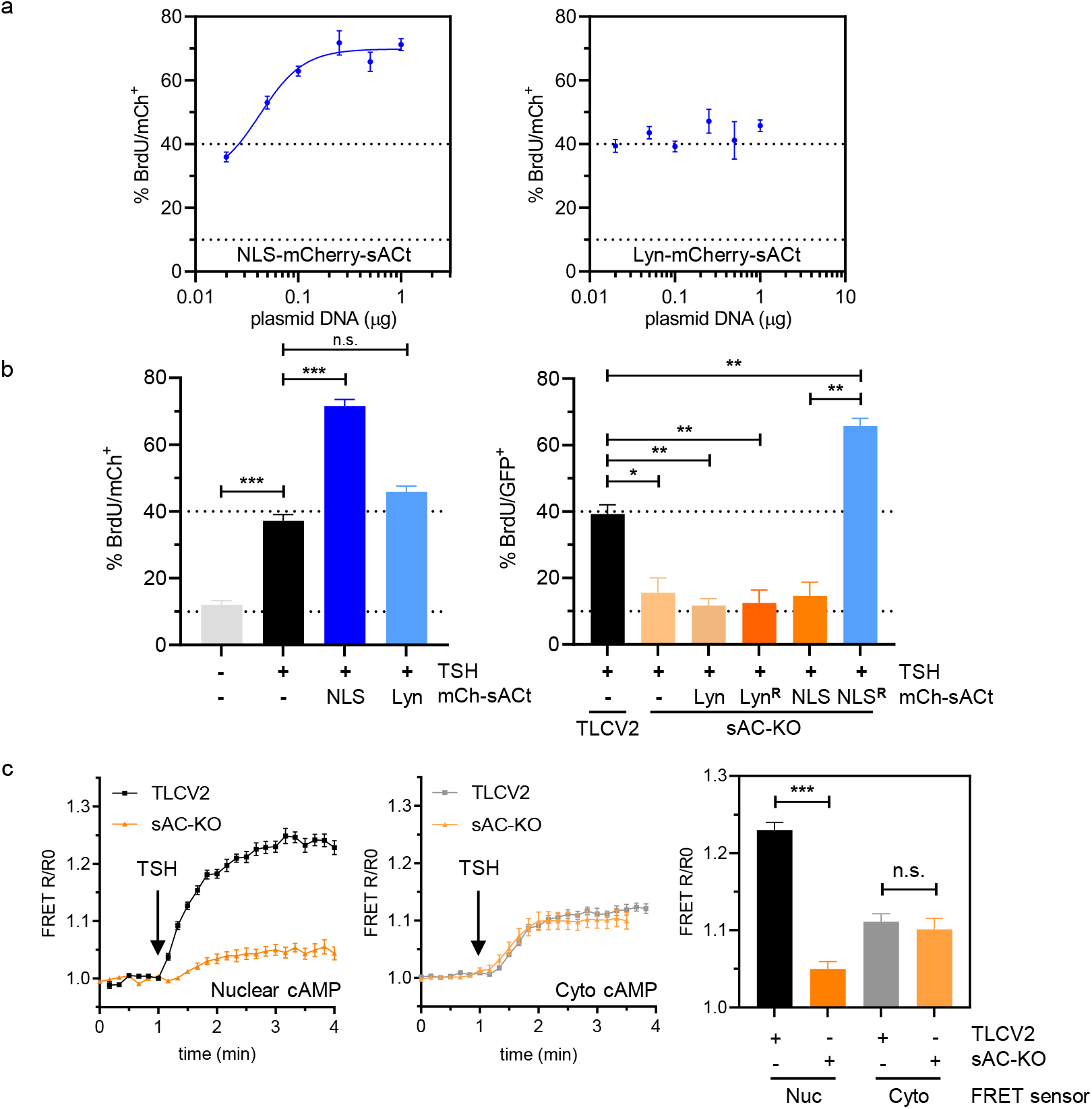
Nuclear-targeted sACt stimulates proliferation in a dose-dependent manner. (a) PCCL3 cells were transfected with increasing doses of the nuclear-targeted NLS-mCherry-sACt (left panel) or the membrane-targeted Lyn-mCherry-sACt constructs (right panel) in the presence of TSH (1 mIU/ml). Proliferation was assessed using BrdU incorporation. (b) Proliferation was assessed in the absence or presence of TSH and expression of NLS-mCherry-sACt or Lyn-mCherry-sACt plasmids (left panel). CRISPR/Cas9-mediated sAC deletion using the TLCV2-sg243 but not the TLCV2-vector successfully inhibited cell proliferation assessed by BrdU incorporation. Proliferation was successfully rescued by an sg-resistant NLS-mCherry-sACt^R^. but not by the sg-sensitive NLS-mCherry-sACt, Lyn-mCherry-sACt, or Lyn-mCherry-sACt^R^ (right panel). (c) Real-time FRET-based monitoring of cAMP levels in live PCCL3 cells. Traces are the normalized FRET ratios (R/R0) of the NLS-H208 (nuclear) and H208 (cytosolic) sensors. Nuclear cAMP measurements were performed in cells transfected with TLCV2-vector (TLCV2) or TLCV2-sg243 (sAC-KO) using the nuclear (left panel) or the cytosolic cAMP sensors (middle panel). Quantitative analysis of R/R0 values after 3 min of TSH stimulation in the nuclear and cytosolic compartments (right panel). BrdU data are expressed as % BrdU/mCh^+^ or GFP^+^ cells. Data are expressed as mean ± SEM (for n cells/samples and independent experiments, see statistics). Significance was tested using a two-tailed Student’s t-test (*, p<0.05; **, p<0.01; ***, p<0.001; n.s., not significant).

### sAC inhibition cannot be rescued by a membrane-permeable cAMP analog

8-Bromoadenosine 3’,5’-cyclic monophosphate (8-Br-cAMP) is a membrane-permeable cAMP analog capable of activating both Epac1 and PKA ^58^, cAMP effectors involved in the TSH mitogenic response ^19^. Accordingly, 8-Br-cAMP mimicked TSH and was sufficient to trigger proliferation in PCCL3 cells assessed by both ^3^H-thymidine and BrdU incorporation assays (Fig. 4a). Although we initially reasoned that 8-Br-cAMP, mimicking the cyclase reaction product, should bypass the negative effects of sAC inhibition, preincubation of cells with LRE1 or KH7 consistently reduced cell proliferation. (Fig. 4b). Although 8-Br-cAMP incubation was able to raise nuclear cAMP levels, this effect remained sensitive to LRE1. Incubation with FK and IBMX further increases nuclear cAMP in an LRE1-resistant manner (Fig. 4c). Similarly, in HC-1 cells that show very low basal cAMP levels ^55^, incubation with 8-Br-cAMP manifests a slow increase in the cytosol that is absent in the nucleus for at least 30min (Figs. S7a-b). However, upon AC9 transfection, that in the absence of stimulation does not increase cAMP levels, 8-Br-cAMP incubation dramatically increases the rate of cytosolic cAMP accumulation with a kinetic profile consistent with the sustained cAMP wave, revealing now a nuclear LRE1-sensitive accumulation (Fig. S7c-d). Therefore, these combined results show that 8-Br-cAMP itself is unable to reach the nuclear compartment even at concentrations where it effectively stimulates cell proliferation. This indicates that its effects, like those of TSH, require a sustained cAMP-dependent activation of nuclear sAC, an enzyme strictly required for cell proliferation.

**Fig. 4:**
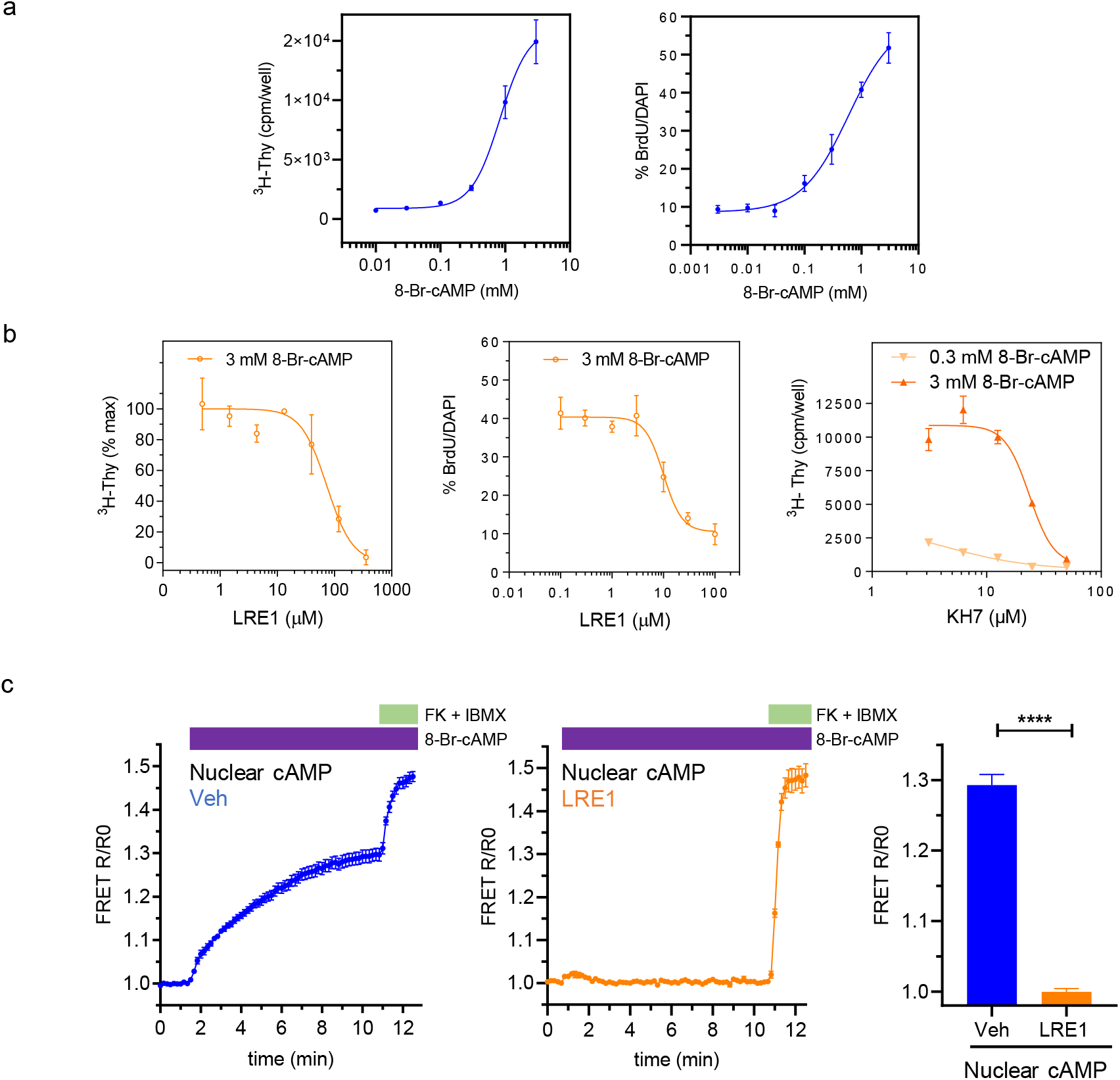
8-Br-cAMP requires an active sAC to stimulate proliferation. (a) Proliferation was assessed by ^3^H-thymidine (left panel) and BrdU incorporation (right panel) in PCCL3 cells incubated with increasing doses of 8-Br-cAMP. (b) Cells were incubated with 3 mM 8-Br-cAMP and increasing doses of LRE1. Proliferation was assessed by ^3^H-thymidine (left panel) and BrdU incorporation (middle panel). Cells were incubated with 0.3 or 3 mM 8-Br-cAMP and increasing doses of KH7 (right panel). ^3^H-thymidine data are expressed as the averaged counts per minute in each well (cpm/well). Curve fitting was performed in GraphPad Prism. (c) Real-time FRET-based monitoring of cAMP levels in live PCCL3 cells. Traces are the normalized FRET ratios (R/R0) of the NLS-H208 (nuclear) cAMP sensor. Normalization was determined by averaging 3 timepoints before the addition of 8-Br-cAMP. cAMP measurements were performed in cells preincubated with Vehicle (Veh; DMSO, left panel) or LRE1 (100 μM; middle panel). Quantitative analysis of R/R0 values after 5 min of TSH stimulation in the nuclear compartment (right panel). Cells were stimulated first with 8-Br-cAMP and then successively with FK (20 μM) and IBMX (250 μM). Data are expressed as mean ± SEM (for n cells/samples and independent experiments, see statistics). Significance was tested using a two-tailed Student’s t-test (****, p<0.0001).

### Nuclear sAC is required for local PKA activation and CREB phosphorylation

PKA and its substrate, CREB are the immediate downstream players in the cAMP pathway required for cell proliferation of PCCL3 cells ^19^. To evaluate if sAC is the source of cAMP activating these effectors we first assessed if nuclear PKA activity increases after TSHR activation. In cells transfected with NLS-AKAR4, a nuclear-targeted FRET-based PKA activity sensor ^45^, TSH incubation elicited a rapid increase in nuclear PKA activity, with a fast initial peak consistent with the nuclear cAMP kinetics, followed by a slower sustained phase (Figs. 5a, S8a). Both pharmacological and down-regulation experiments performed with LRE1 and sh-sAC respectively showed inhibition of local PKA activity, mainly affecting the fast initial peak (Fig. 5a-c). To evaluate CREB phosphorylation cells were starved for 20 h in a TSH-free and reduced FBS (0.5%) medium and then stimulated with TSH at saturating concentrations. pCREB levels were evaluated at different times by immunofluorescence and immunoblotting, showing a positive signal in most cells (~80%) after 10 minutes of TSH incubation (Fig. 5d, S8b-c). Pharmacological (LRE1) and genetic (sh-sAC) inhibition of sAC impeded CREB phosphorylation (Fig. 5d-f, S8b-d). These results show that nuclear sAC activity is essential for the activation of two successive elements of the cAMP pathway, PKA and CREB. In sum, the evidence presented so far strongly suggests that the levels of cAMP in the nuclear compartment determine G1/S progression.

**Fig. 5:**
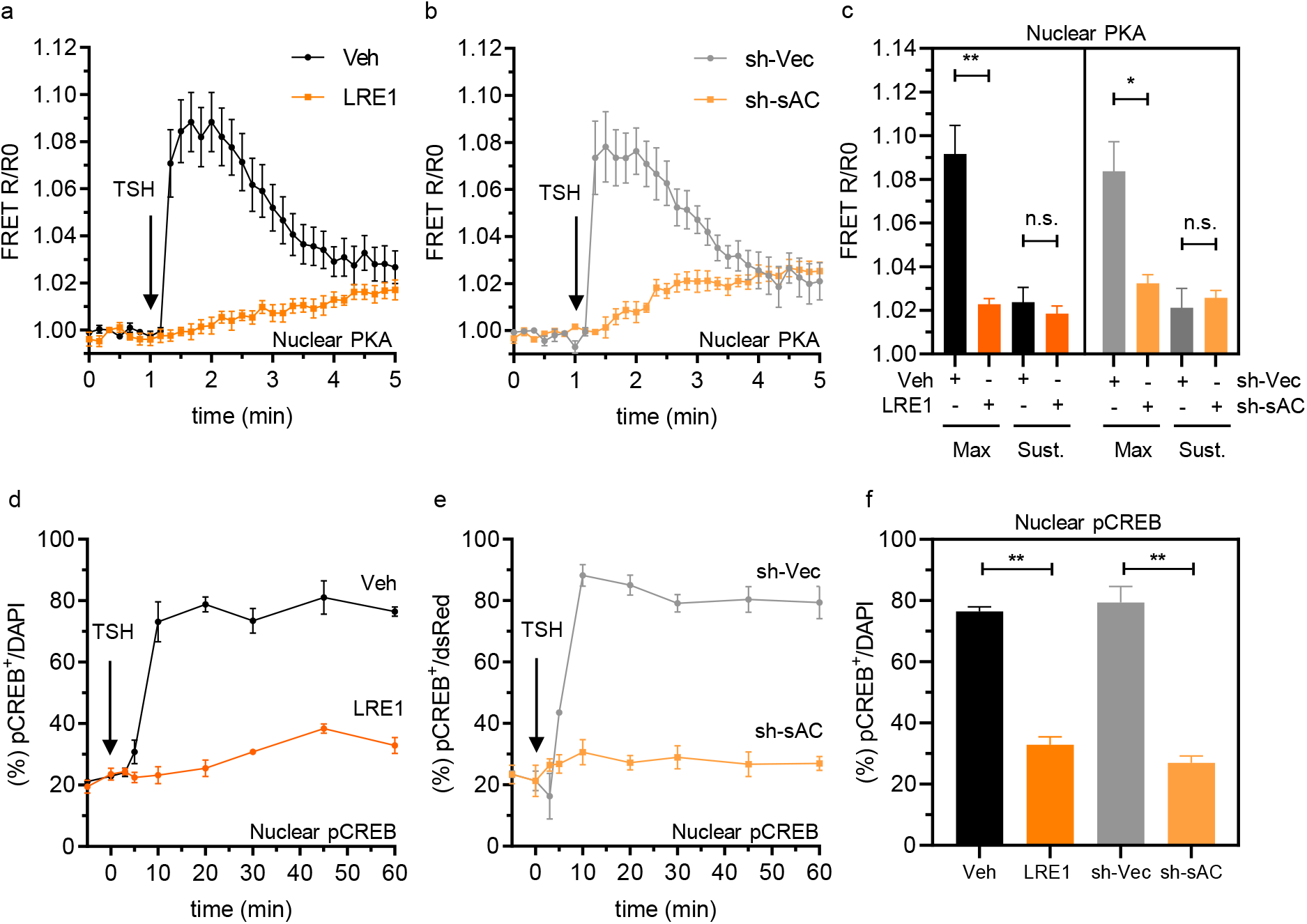
Nuclear PKA and pCREB are activated in a sAC-sensitive manner. Real-time FRET-based monitoring of PKA activity in live PCCL3 cells. Traces are the normalized FRET ratios (R/R0) of the NLS-AKAR4 sensor. Normalization was determined by averaging 3 timepoints before the addition of TSH (1 mIU/ml). (a) PKA activity measurements were performed in cells preincubated with either Vehicle (Veh; DMSO) or LRE1 (100 μM) 30 min before TSH stimulation. (b) PKA activity measurements were performed in cells transfected with either sh-sAC or sh-Vec and stimulated with TSH. (c) Quantitative analysis of either maximum (Max) or sustained (Sust.) R/R0 values after TSH stimulation in the nuclear compartment. Max: the maximum R/R0 values during the initial 1 min of TSH stimulation were considered; Sust.: R/R0 values after 5 min of TSH stimulation were considered. (d) CREB phosphorylation was assessed by immunofluorescence at 0, 3, 5, 10, 20, 30, 45, and 60 min after TSH stimulation. Data are the % of pCREB^+^/DAPI at each timepoint. Cells were preincubated with Vehicle (Veh, DMSO) or LRE (100 μM). (e) CREB phosphorylation was assessed in cells transfected with either sh-Vec or sh-sAC. (f) Quantitative analysis of nuclear pCREB^+^/DAPI (%) values after 60 min of TSH stimulation. Data are the % of pCREB^+^/dsRed^+^ cells at each timepoint. Data are expressed as mean ± SEM (for n cells/samples and independent experiments, see statistics). Significance was tested using a two-tailed Student’s t-test (*, p<0.05; **, p<0.01; n.s. not significant).

### Nuclear but not cytosolic cAMP levels control G1/S progression

To assess whether nuclear cAMP levels alone can control cell cycle progression, we exploited bPAC-nLuc, an optogenetic tool recently developed in our laboratory ^50^,^59^, that combines a blue light-activated (445 nm) adenylyl cyclase (bPAC) and luciferase (nLuc). Stably transfected NLS-bPAC-nLuc cells responded to TSH as well as to furimazine and blue light stimulation (see Methods). Compared to TSH, and like our results with the untargeted and NLS-sACt constructs (Figs. 2b and 3b), proliferation rose to 60% in cells stimulated with furimazine or blue light (Fig. 6a). Although preincubation of cells with LRE1 consistently blocked proliferation triggered by NLS-sACt, the sAC inhibitor did not affect NLS-bPAC-nLuc stimulation (Fig. 6b). Moreover, a single blue light pulse (0.5 s) triggered CREB phosphorylation but, unlike TSH, it was insensitive to LRE1 (Fig. S9a-b). These results demonstrated that NLS-bPAC-nLuc-mediated nuclear cAMP generation is sufficient for proliferation in an LRE1-insensitive manner. However, even though we confirmed that NLS-bPAC-nLuc expression is restricted to the nuclear compartment and that it generates cAMP signals in the nucleus, we also found a minor cAMP ‘leak’ towards the cytosol (Fig. S10b) ^59^. To rule out that this leak was responsible for activating one or more cytosolic effectors, we utilized ΔRI-PDE8, a PDE8-derived construct with an increased hydrolytic activity ^60^ that does not bind to AKAPs (Fig. S9d) (see Methods). We reasoned that targeting ΔRI-PDE8 to the ER membrane facing the cytosol with a P450 sequence ^61^, will rapidly inactivate any cAMP diffusing out of the nucleus into the cytosol. Both the nuclear (NLS) and the ER-cytosolic (P450) targeted versions of ΔRI-PDE8 were successful in specifically blocking cAMP local elevations without affecting the other compartment response (Fig. S9c). In addition, to rule out that the cAMP ‘leak’ was not actually cytosolic NLS-bPAC-nLuc expression, cells were photostimulated with a 445 nm laser controlled by a point scanning system that allowed us to stimulate a small area of ~1 μm^2^ within the cytoplasm (Fig. S9e) (see Methods). While laser pulses directed to the nuclear area were able to generate cAMP elevations (detected both in the nuclear and cytosolic compartments), pulses directed to the cytosol of the same cells were not (Fig. S10a-b). In this context, NLS- and P450-ΔRI-PDE8 expression successfully abolished nuclear and cytosolic cAMP elevations generated by nuclear-directed light pulses (Fig. S10c-e). Finally, while both ΔRI-PDE8 constructs were able to reduce proliferation in WT-PCCL3 cells, light-stimulated NLS-bPAC-nLuc expressing cells proliferated in the presence of P450-ΔRI-PDE8 but not in the presence of NLS-ΔRI-PDE8 (Fig. 6c-d). These results conclusively demonstrate that nuclear but not cytosolic cAMP is the key factor defining cell cycle progression.

**Fig. 6:**
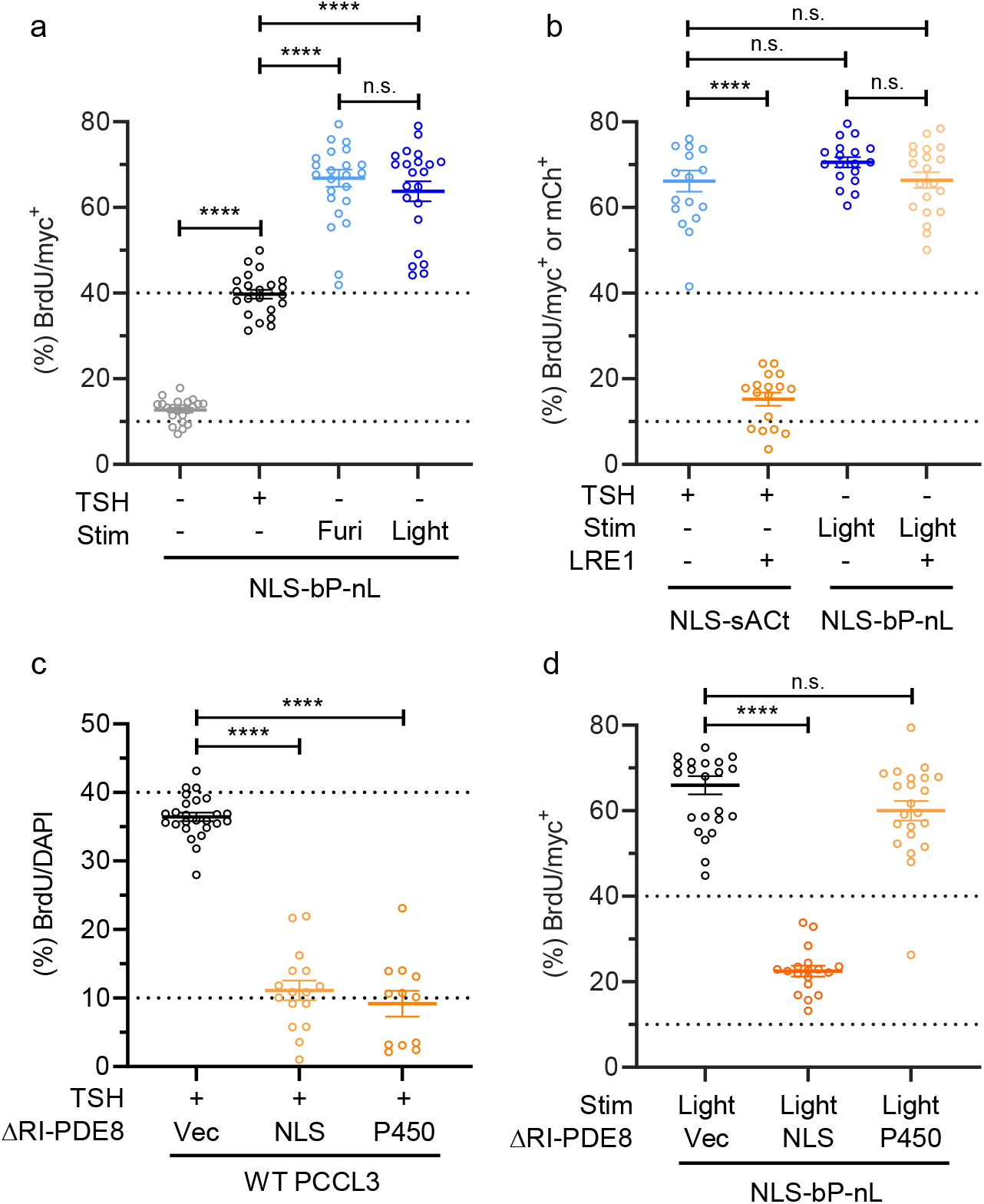
Nuclear cAMP is sufficient for cell proliferation. (a) PCCL3 cells were stably transfected with NLS-bPAC-nLuc and proliferation was assessed using BrdU incorporation. In the absence of light stimulation proliferation was TSH-dependent (1 mIU/ml). In the absence of TSH, cells proliferated when exposed to chemical stimulation (Furimazine, Fz-4377; Furi, 1:1000) or light (440 nm, 0.5s, 15 min interval) stimulation for 24 h. (b) Cells were transiently transfected with NLS-mCherry-sACt and stimulated with TSH or stably transfected with NLS-bPAC-nLuc and stimulated with light (440 nm, 0.5s, 15 min interval) in the absence of TSH. Proliferation was assessed in the presence of Vehicle (Veh; DMSO) or LRE1 (100 μM). (c) WT-PCCL3 cells were transfected with the NLS-ΔRI-PDE8, P450-ΔRI-PDE8 or empty vector (Vec), and proliferation was assessed in the presence of TSH. PCCL3 cells stably transfected with NLS-bPAC-nLuc were transiently transfected with an empty vector (Vec), NLS-ΔRI-PDE8, or P450-ΔRI-PDE8 constructs and stimulated with light (440 nm, 0.5s, 15 min interval). BrdU data are expressed as % BrdU/mCh^+^ or GFP^+^ cells. Data are expressed as mean ± SEM (for n cells/samples and independent experiments, see statistics). Significance was tested using a two-tailed Student’s t-test (****, p<0.0001; n.s. not significant).

## DISCUSSION

Using a combination of pharmacological, genetic, and optogenetic approaches we show that agonist-dependent TSHR-Gs-tmAC activation at the plasma membrane, in an internalization-dependent manner, results in fast nuclear sAC-mediated cAMP production, PKA activation, and CREB phosphorylation, critical for maintaining cell proliferation. We also show that small variations in the nuclear expression levels of sAC have a large impact on the proliferative response, consistent with nuclear sAC activation representing a rate-limiting step. Furthermore, our findings show that an elevation of cAMP in the nucleus, independent of a cytosolic cAMP component, is sufficient to stimulate cell proliferation.

For many years, PKA-C translocation to the nucleus upon cAMP-mediated activation was thought to represent the rate-limiting step for subsequent nuclear transcriptional events. A diffusive mechanism was proposed ^62^ and recently estimated to represent ~1–2% of total cellular PKA-C ^40^. However, its slower kinetics compared to nuclear cAMP accumulation and PKA activation rates suggested that cAMP diffusion into the nucleus rather than PKA-C translocation might represent the rate-limiting step, acting instead on a local nuclear-resident PKA pool ^39^. However, our 8-Br-cAMP results indicate that despite its small molecular weight and diffusivity, cAMP is not itself accumulating in the nucleus, it is most likely restricted in the cytosol by binding and/or PDE activity. Instead, we propose that cAMP, synthesized intracellularly, is rather required to generate an intermediate regulator responsible for an LRE1-sensitive nuclear sAC activation. Only upon non-physiological strong stimulation with FK and/or IBMX, cAMP concentration overcomes the buffering capacity and/or PDE nanodomains, allowing it to accumulate in the nucleus in an LRE1-insensitive manner.

Upon activation, nuclear PKA activity is terminated by binding to PKI, resulting in PKA-C inhibition and nuclear export ^63^. TSH-mediated nuclear PKA activation showed a fast sAC-dependent initial phase, followed by a delayed sAC-independent sustained phase (Figs. 5a-c). It is tempting to speculate, that this sustained phase might represent the slow translocation of PKA-C as a mechanism to replenish the local nuclear PKA pool. This testable hypothesis deserves further investigation.

In summary, our combined results utilizing sAC inhibition/downregulation and the effects of 8-Br-cAMP clearly indicate that neither PKA-C translocation nor cAMP diffusion, but rather nuclear sAC activation is the rate-limiting step responsible for the fast nuclear cAMP accumulation, PKA activation, and CREB phosphorylation.

Recent works have shown that sAC may be an alternative cAMP source amplifying the GsPCR signaling in many cell types via Ca^2+^ and/or HCO_3_^-^. Prostaglandins via EP1 and EP4 receptors in bronchial cells can activate sAC in a mechanism that depends on Ca^2+^ but not HCO_3_^-^ ^64^,^65^. Corticotropin-releasing hormone receptor 1 (CRHR1) in hippocampal neurons can activate sAC in an internalization-dependent manner, generating a sustained cAMP signal; CRH-mediated sAC activation depends on both Ca^2+^ and HCO_3_^-^ ^66^. Follicle-stimulating hormone receptor (FSHR) activation in the granulosa cells in the ovary stimulates sAC in a mechanism that depends on Cystic Fibrosis Transmembrane Regulator (CFTR)-mediated HCO_3_^-^ influx ^67^. Similarly, Luteinizing hormone receptor (LHR) stimulation in testicular Leydig and Sertoli cells activated sAC, in a mechanism proposed to be dependent on CFTR/HCO3^-^ ^68^, consistent with the involvement of a CFTR/HCO3^-^/sAC-dependent cAMP signalling in spermatogenesis ^69^. Interestingly, Epac-Rap has been linked to both CFTR ^64^,^70–72^ and Ca2^+^ influx and mobilization from intracellular stores (reviewed in ^73^). We propose that an internalization-dependent activator of sAC either diffuses into the nucleus or is produced locally, triggering nuclear events (*e.g*., PKA activation, CREB phosphorylation, and transcriptional activation) involved in cell proliferation (Fig. 7). Future works should address if HCO_3_^-^ and/or Ca^2+^ represent this unknown intermediate diffusible factor that drives proliferation into thyroid cells and test the intriguing possibility that the functional role of the intracellular sustained cAMP wave is the generation of this intermediate required for nuclear sAC activation.

**Fig. 7:**
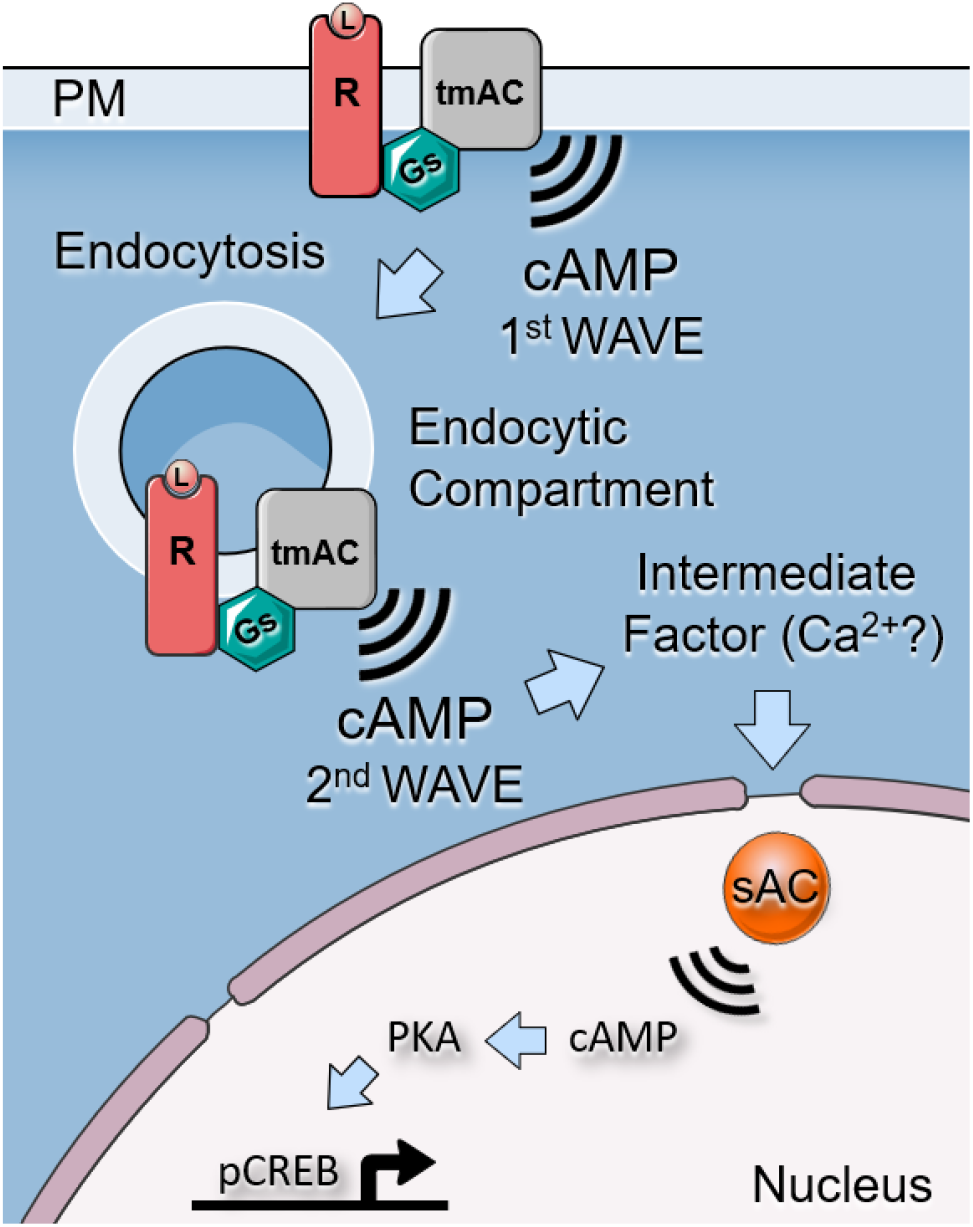
Soluble cyclase-mediated nuclear cAMP synthesis is required and sufficient for cell proliferation. Ligand-mediated GPCR activation triggers a Gαs-mediated cAMP synthesis by one or more tmACs generating the first cAMP wave in the plasma membrane compartment. Ligand binding also triggers the internalization of the signaling complex that drives cAMP production into the endocytic compartment. Our combined results suggest that cAMP generated in the endocytic compartment (second wave) facilitates the production of an intermediate factor that, unlike cytosolic cAMP, can reach the nuclear compartment and rapidly activate nuclear sAC. Local cAMP production quickly activates a local PKA pool which leads to CREB phosphorylation activating transcription of cAMP-dependent genes and enhancing cell proliferation. Our results also suggest that proliferation may also be sustained solely by the elevation of nuclear cAMP, independent of the events occurring in the membrane or the cytosolic compartments.

Finally, in this work we also demonstrated that cAMP generated in the nucleus is sufficient to trigger cell proliferation. Utilizing novel compartment-specific cAMP actuators to synthesize (NLS-bPAC-nLuc) or hydrolyze (ΔRI-PDE8) cAMP, we show that light-dependent nuclear cAMP synthesis is sufficient for CREB phosphorylation and G1/S entry in a sAC-independent manner. Moreover, while the expression of a cytosolic ΔRI-PDE8 blocked TSH-mediated cell proliferation, it was unable to affect light-dependent proliferation triggered by NLS-bPAC-nLuc, thus ruling out concerns of a putative cAMP ‘leak’ activating cytosolic effectors and stimulating proliferation. Therefore, we propose that a nuclear PKA pool is rapidly activated downstream of nuclear sAC activation, thanks to local production of cAMP and without any diffusion of cAMP in the nucleus. This leads to phosphorylation of CREB and is sufficient to maintain cell proliferation.

## METHODS

### Materials

Forskolin (F6886), IBMX (I7018), LRE1 (SML1857), 8-Br-cAMP (B7880), BrdU (B5002), Dynasore (D7693), (-)-Isoproterenol (I6504) and Dowex-Alumina resins were from Sigma. Dyngo-4a (D4a; ab120689) was from Abcam. [2,8-^3^H]-Adenine (NET063001MC) was from Perkin Elmer. TSH (TSH-1315B) was from Creative Biomart. Furimazine (Fz-4377) was from Promega. KH7 and inactive analog KH7.15 were provided by Drs Buck-Levin (Cornell).

### Antibodies

#### Primary

Anti-myc (9E10); Covance MMS-150R

Anti-c-Myc-FITC; Sigma F2047

Anti-c-Myc-Cy3; Sigma C6594

Anti-CREB (9104) and anti-pCREB (9198s); Cell Signaling

Anti-FLAG; Sigma F1804

DyLight-650 Anti-FLAG; Abcam ab117492

Anti-FLAG-Cy3; Sigma A9594

Anti-FLAG-FITC; Sigma F4049

Sheep anti-BrdU; Capralogics; P00013

Anti-sAC (ADCY10); Thermo PA5-25756

Anti-GFP; Invitrogen A-11122 Anti-Actin; Sigma A5316

#### Secondary

Donkey Anti-Sheep Alexa-647; Abcam ab150179

Donkey Anti-Sheep Alexa-488; Thermo A-11015

Donkey Anti-Mouse Alexa-488; Thermo A-21202

Donkey Anti-Mouse Alexa-649; Thermo A10038

Donkey Anti-Rabbit Alexa-488; Thermo A-21206

Goat Anti-Mouse Alexa-555; Thermo A32727

Goat Anti-Rabbit Alexa-594; Thermo A32740

Goat Anti-Rabbit Alexa-647; Thermo A-21244

### DNA constructs

The cAMP FRET sensors H188 and H208 (YFP-EPAC-Q270E-mScarletI) were kindly provided by Dr. Jalink and were reported before ^50^. NLS-H188 and NLS-H208 were constructed using PCR to subclone an SV40-NLS motif to the N-terminal by using HindIII. NLS-AKAR4 (Plasmid #138217), pcDNA3-sACt-mCherry-NLS (Plasmid #138214), pcDNA3-Lyn-sACt-mCherry (Plasmid #138216) and HA-Dyn1-K44A (Plasmid #34683) plasmids were obtained from Addgene. The NLS-ΔRI-PDE8 and P450-ΔRI-PDE8 plasmids contain a deletion of aa1-74 in RI from the original template ^60^, and were generated by the addition of 3xSV40 NLS-FLAG and P450 2C1 ER ^61^ targeting-FLAG to their N-termini, respectively. Expression was assessed with anti-FLAG antibodies. pcDNA3.0 AC2-HA was a gift from Dr. Cooper (Cambridge University). Flag-hAC9 was kindly provided by Lily Jiang (UT Southwestern). The shRNA-sAC(1-2), sh-scramble (sh-Vec), and expression plasmids for sACt and sACt^R^ were described in ^74^ and provided by Dr. Muller (Northwestern University). Sh-Vec and sh-sAC were transferred to pSIREN-DNR-dsRedExpress. sACt and sACt^R^ were transferred to pCMV-myc.

### Cell lines and transfections

PCCL3, a normal TSH-dependent rat thyroid follicular cell line, was grown in Coon’s Media: Nutrient Mixture F-12 Ham (Coon’s modification; Sigma-Aldrich, F6636) supplemented with 2.68 g/L sodium bicarbonate, 5% fetal bovine serum (FBS; Corning, MT35011CV), 1% Penicillin/Streptomycin, 2 mM L-glutamine, insulin (1 μg/mL), apo-transferrin (5 μg/mL), hydrocortisone (1 nM), and thyroid-stimulating hormone (TSH 1 mIU/ml). The rat hepatoma clonal cell line HC1 (kindly provided by Dr. Elliot Ross, University of Texas Southwestern Medical Center) was grown in DMEM (Corning, 10013CV) supplemented with 10% FBS, penicillin (100 IU/L), and streptomycin (100 mg/L), and L-glutamine. Cells were grown to ~90% confluency before passaging every 2-4 days at 37°C in 5% CO_2_, 95% humidified air. Transient transfections were performed using Lipofectamine 3000 Transfection kit (Invitrogen) for 24h. NLS-bPAC-nLuc stable cell lines were generated by lentiviral infection with a multiplicity of infection (MOI) of 80 and 5 μg/mL of polybrene and selected with puromycin as described before ^50^.

### Generation of sAC KO cells

sAC-KO PCCL3 cells were generated by CRISPR/Cas9 genome editing system. Several sg sequences were identified with CHOPCHOP (https://chopchop.cbu.uib.no/). sgRNAs were synthesized and subcloned into the BsmBI site of the all-in-one (dox-inducible Cas9-2A-eGFP and constitutive U6 promoter) TLCV2 vector (Addgene #87360). sAC-targeted sgRNAs: sg243 to exon 2 of C1 domain and sg396 to exon4 of C1 domain (see below).

243 sense 5’-caccgCTACTACATAAGTGCGATAG-3’
243 anti-sense 5’-aaacCTATCGCACTTATGTAGTAGc-3’
396 sense 5’-caccgTGGCTTGTTTGAAGCCAAGG-3’
396 anti-sense 5’-aaacCCTTGGCTTCAAACAAGCCAc-3’

For lentivirus production, HEK293T cells were seeded on 10 cm tissue culture dishes and incubated until cells reached ~70% confluence. TLCV2, TLCV2-sg243, or TLCV2-sg396 constructs were mixed with packaging vectors pMD2.G and pCMV-delta-R8.2 and transfected using X-tremeGENE HP (Roche). The virus-containing medium was collected and filtered. Lentivirus was concentrated using the Lenti-X concentrator (TaKaRa). Cells were selected for six days in puromycin followed by three days induction with doxycycline. Gene editing efficiency in the puro-resistant clones was assessed by a T7e assay (CRISPR Genomic Cleavage Detection Kit, Abm). The sg243-resistant sACt-mCherry plasmids were made by introducing the following (underlined) mutations (CTA<u>T</u>TATAT<u>CTCC</u>GC<u>C</u>ATCGT<u>C</u>).

### FRET-based cAMP measurements

Cells were seeded on 25 mm glass coverslips and transfected with the H188, H208, NLS-H208, or NLS-AKAR4 FRET-based sensors. Transfected cells were given fresh media for 48 h and hormone starved for 3 h in Coon’s media with 5% FBS and lacking TSH, insulin, and hydrocortisone before measurements. Cells were washed once in PBS and imaged in OptiMEM media lacking phenol red (Gibco). Two microscope setups were used; Setup A: Olympus IX70 microscope equipped with a Till Polychrome V monochromator, a 60x/1.4 NA oil objective, and a Hamamatsu CCD camera (Photonics Model C4742-80-12AG; 8×8 binning). Setup B: Olympus IX83 Motorized two-deck Microscope equipped with a 6-line multi-LED Lumencor Spectra X, a Prior Emission Filter Wheel, Prior Proscan XY stage, an ORCA-Fusion Digital CMOS camera (C14440-20UP; 4×4 binning), a 40x/1.4 NA oil objective (UPLXAPO30X). Images were acquired every 5 or 10s depending on the experiment, using Slidebook 6 software (Intelligent Imaging Innovations Inc.).

The NLS-AKAR and H188 sensors sensor were excited at 440nm and fluorescence was collected using emission filters 470/30nm and 535/30nm with a 69008bs dichroic (Chroma Technology Corp.). The H208 and NLS-H208 sensors were excited at 500nm with emission filters 510/20 and 620/52 and two dichroic beam splitters: 468/526/596 (Semrock) or 69008bs (Chroma Technology Corp) for setups A and B respectively. No significant photobleaching was observed during the time-lapses. Changes in background-subtracted FRET ratios (R) relative to resting conditions (R/R0) were calculated using 3 timepoints before the addition of TSH. Statistical analyses were performed using GraphPad Prism 9 (GraphPad Software Inc.) and significance was tested using a two-tailed *Student’s* t-test (α level was defined as 0.05).

### Optogenetic Stimulation

Cell culture and experiments with cells expressing NLS-bPac-nLuc were performed in a dark environment using a red safelight lamp (Kodak GBX-2 Safelight Filter) with a 13W amber compact fluorescence bulb (Low Blue Lights, Photonic Developments LLC) to prevent light exposure from wavelengths <500nm. Light activation was achieved using a custom-built, Arduino-controlled system capable of regulating the duration, frequency, and intensity of light exposure ^50^,^59^. The illuminating high-power LED (royal blue CREE XTE Tri-Star LED, LED Supply) was mounted on a stage, keeping a distance of 18cm from the sample to ensure a near-homogenous spread of light. For experiments performed inside an incubator, samples were placed atop the stage, and cells were illuminated from below. The stage was inverted for FRET-based experiments, such that cells were illuminated from above. The light intensity used for these experiments was 4.41 ± 0.30 μW/mm2 as measured by a laser power meter (ThorLabs PM1100D, detector S130C). In some experiments, cells were stimulated with a solid-state laser illuminator (445nm, LDI-7, 89 North) connected to a UGA-42 Geo (Rapp OptoElectronic) point scanning system that allowed us to simulate a very small circular area (~1 μm^2^) inside the nucleus (see Fig. S9e). This device was coupled into one of the backports of a two-deck microscope (setup B) allowing us to stimulate cells and perform FRET measurements simultaneously by using an independent lightpath with a ZT458rdc dichroic (Chroma Technology Corp.). While the photobleaching test was performed with a 10 % physical ND filter in the laser light path (see Fig. S9e), all the experiments with live cells were made with a 1% ND.

### Adenylyl cyclase activity in cells

Labeling of cellular ATP pool was performed by overnight incubation with 1mCi/ml [^3^H]-adenine in complete Coon’s medium. The next day cells were washed twice and incubated for 1h in Coon’s starvation medium. Cells were then stimulated with TSH together with either vehicle or inhibitors. Reactions were stopped in trichloroacetic acid (7.5% w/v). Product [^3^H]-cAMP was separated from the substrate [^3^H]-ATP by sequential column chromatography over Dowex and Alumina, as previously described ^75^.

### Immunofluorescence and BrdU Labeling

Cells were grown to 70% confluency on glass coverslips, incubated with Lipofectamine 3000 Transfection kit (Invitrogen) and the corresponding plasmids for 24 h Then after 24-48 h cells were made quiescent by TSH starvation in Coon’s (5% FBS) media for 16 h Upon agonist stimulation (TSH; 1 mUI/ml) for 8 h, cells were labeled for 16 h with BrdU (Sigma, 100 μM). At the end of the labeling period, incorporated BrdU was detected by indirect immunofluorescence (IF). Unless otherwise noted, IF protocol started with fixation in 4% paraformaldehyde (10 min, room temperature) and permeabilization with 0.5% Triton X-100 (10 min, room temperature). After washing, cells were stained for 30 min at 37°C with the primary antibody diluted in PBS/3% BSA/0.05% Tween 20 (see antibodies section). After washing, samples were incubated for 30 min at 37°C with the secondary antibody diluted in PBS/3% BSA/0.05% Tween 20 (see antibodies section) and 4’,6-diamidino-2-phenylindole (DAPI 0.125 mg/ml, Invitrogen) in PBS/3% BSA/0.05% Tween 20. After extensive washes, samples were mounted in Vectashield Vibrance Mounting Medium (Vector Labs; H-1700-10). For BrdU experiments, a sheep anti-BrdU antibody was used in the presence of RQ1 DNase (Promega, 10 units/ml). and after washing with conjugated anti-sheep antibodies (see antibodies section). Data are expressed as the % of BrdU+ cells among the total population (DAPI) or a particular population of cells expressing different fluorescent (dsRed, mCherry, GFP) or non-fluorescent (myc, FLAG) tags. Significance was tested using a two-tailed *Student’s* t-test (α level was defined as 0.05). The immunofluorescence images were acquired in setups A or B described above, or in an Olympus Fluoview FV1000 confocal imaging system.

### ^3^H thymidine proliferation assay

Thymidine incorporation assay was performed as described before ^51^. Briefly, cells were plated into 96-well plates (10,000 cells/well). On the next day, cells were made quiescent by serum starvation in Dulbecco’s modified Eagle’s medium, 0.2% BSA for 20h. Upon agonist stimulation (16h), cells were labeled with [methyl-^3^H]-thymidine (Amersham Biosciences; 1 μM, 1 μCi/ml), and 24h later, samples were collected by using a cell harvester. Filters were dried and analyzed by scintillation counting.

### Statistics

Comparisons were made using an unpaired Student’s t-test with Welch’s correction of n cells/samples and N independent experiments (n/N) as described below. For panels with one or more time points/concentrations, we report the one with the fewest number of independent experiments. Statistical analyzes and curve fitting were performed using Graph Pad Prism 9 (GraphPad Software Inc.). P values less than 0.05 (p < 0.05) were considered significant.

Fig. 1: a (14/6); b (7/4); c: Veh (6/4), D4a (7/3); d: Veh (8/3), Dyna (12/5); e: Veh (6/3), D44a (11/6); f: Veh (6/3), LRE1 (8/4); g: Veh (8/3), LRE1 (6/4); h: Nuc Veh (6/3), Nuc LRE1 (8/4), Cyto Veh (8/3), Cyto LRE1 (6/4); i (5/3); j: Veh- (3/3), Veh+ (3/3), KH7- (3/3), KH7+ (3/3), LRE1- (3/3), LRE1+ (3/3).

Fig. 2: a: left and right middle panels: KH7 (3/3), KH7.15 (3/3), and LRE1 (3/3); right panel: no TSH (11/5), sh-Vec (9/5), sACt (7/4), sh-sAC+sACt (12/5), sh-sAC (9/4), sh-Vec+sACt_R (15/5), sh- sAC+sACt_R (16/6). c: left: Veh (7/4), LRE1 (7/3), center: sh-Vec (4/3), sh-sAC (5/3), right: sACt (3/3), sACt_R (3/3); d: sh-Vec (6/4), sACt (17/6), sh-sAC (12/5); e: sh-Vec (7/5), sACt (18/6), sh-sAC (32/6).

Fig. 3: a: left and right panels (3/3); b: left and right panels (3/3); c: left: TLCV2 (7/5), sAC-KO (9/6), right: TLCV2 (7/4), sAC-KO (14/6).

Fig. 4: a: left and right panels (3/3); b: left, center, and right panels (3/3); c: Veh (26/5), LRE1 (13/4).

Fig 5: a, b, and c: Veh (7/3), LRE1 (8/4), sh-Vec (6/3), sh-sAC (8/3). c, d, and e: Veh (3/3), LRE1 (3/3), sh-Vec (3/3), sh-sAC (3/3).

Fig 6: a: TSH-/Stim- (22/7), TSH+/Stim- (22/6), TSH-/Furi (25/7), Light (22/5); b: NLS-sACt (18/4), NLS-sACt+LRE1 (17/4), bP-nL (18/5), bP-nL+LRE1 (20/4); c: Vec (25/5), NLS-PDE8 (12/6); P450-PDE8 (16/5); d: bP-nL+Vec (25/4), bP-nL+NLS-PDE8 (18/6); bP-nL+P450-PDE8 (22/4).

---

Fig. S1: b: CTRL (6/3), Dyna 10 μM (10/4), Dyna 15 μM (6/3), Dyna 30 μM (4/3), c (4/3); d: Vec (5/4), D44A (3/3); f: Veh (6/4), D4a (7/3), Veh (8/3), Dyna (12/5), Veh (6/3).

Fig. S2: same as Fig. 1h.

Fig. S3: a (6/3); b (6/3); c (5/3); d: CTRL (7/4); LRE1 (8/4).

Fig. S4: a: Veh (4/3), LRE1 (6/3); b (4/3); c: Veh (4/3), Dyna (4/3).

Fig. S5: c: sh-Vec (6/4), sACt (17/5).

Fig. S6: c: TLCV2 (3/3), sg243 (3/3); d: (3/3) for all traces/bars.

Fig. S7: a (10/4); b (6/3); c (14/4); d: Veh (8/3), LRE1 (6/3).

Fig. S8: d: Veh (3/3), LRE1 (3/3).

Fig. S9: a and b: Dark (3/3), Light (3/3), Light+LRE1 (3/3); c: all conditions: (3/3).

Fig. S10: c: left (5/3), right (3/3); e: NLS-H208 (6/4), H208 (6/4).

## Acknowledgments

We thank Drs. Cooper (Cambridge University) and Lily Jiang (UT Southwestern) for the AC2 and Flag-hAC9 expression plasmids, respectively; Dr. Anand (Pennsylvania State University) for the RI-PDE8 construct, Dr. Muller (Northwestern University) for the sh-RNA and sh-resistant sAC plasmids, and Dr. Ross (University of Texas Southwestern Medical Center) for the HC1 cells. Funding was provided by the NIH grants R01 GM099775 and GM130612 to DLA.

## Conflict of interest

The authors declare that they have no conflicts of interest with the contents of this article.

## Notes

### Competing Interest Statement

The authors have declared no competing interest.

