## Supplementary Information for "Soluble cyclase-mediated nuclear cAMP synthesis is sufficient for cell proliferation"

### SUPPLEMENTARY FIGURES

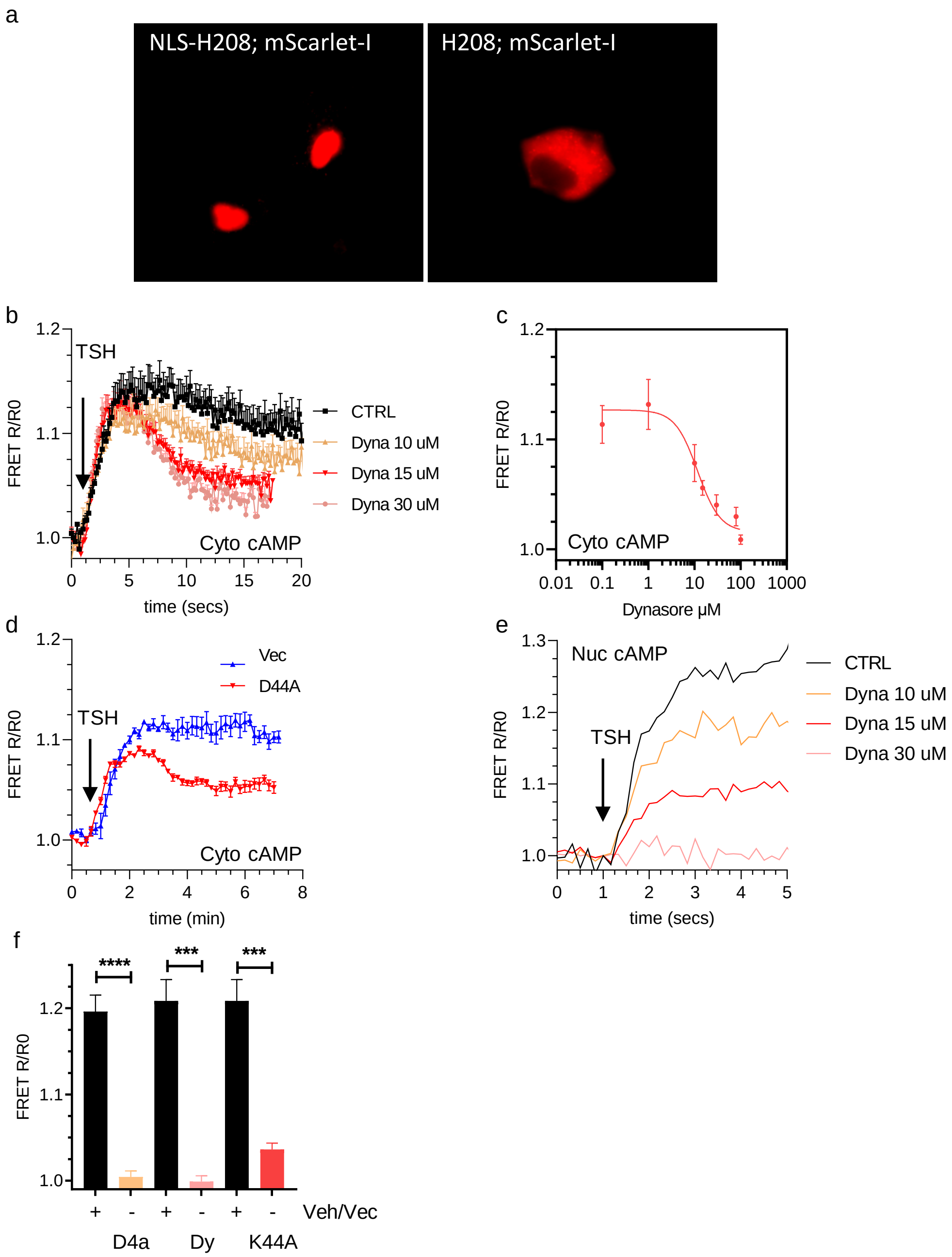


**Fig. S1.** (a) Fluorescence image of NLS-H208 (nuclear; left) and H208 (cytosolic; right) sensors transfected in PCCL3 cells. The fluorescence of mScarlet-I is shown in red. (b) PCCL3 cells were stimulated with TSH (1 mIU/ml). Cytosolic cAMP was measured using the H208 FRET sensor. Dynasore (10, 15, 30 μM was preincubated for 15 min before experiments. (c) Concentration-dependent inhibition of the TSH response by Dynasore. (d) Cells transfected with H208 were cotransfected Vector (Vec) or Dynamin-K44A (D44A). (e) Cells were transfected with NLS-H208 and were stimulated with TSH after 15 min of preincubation with different concentrations of Dynasore. (f) Quantification of the same traces shown in panels c, d and e of Fig. 1 after 2 min of incubation with TSH. Data are expressed as mean ± SEM (for n cells/samples and independent experiments, see statistics). Significance was tested using a two-tailed Student's t-test (***, $p<0.01$; $p<0.001$; n.s., not significant).

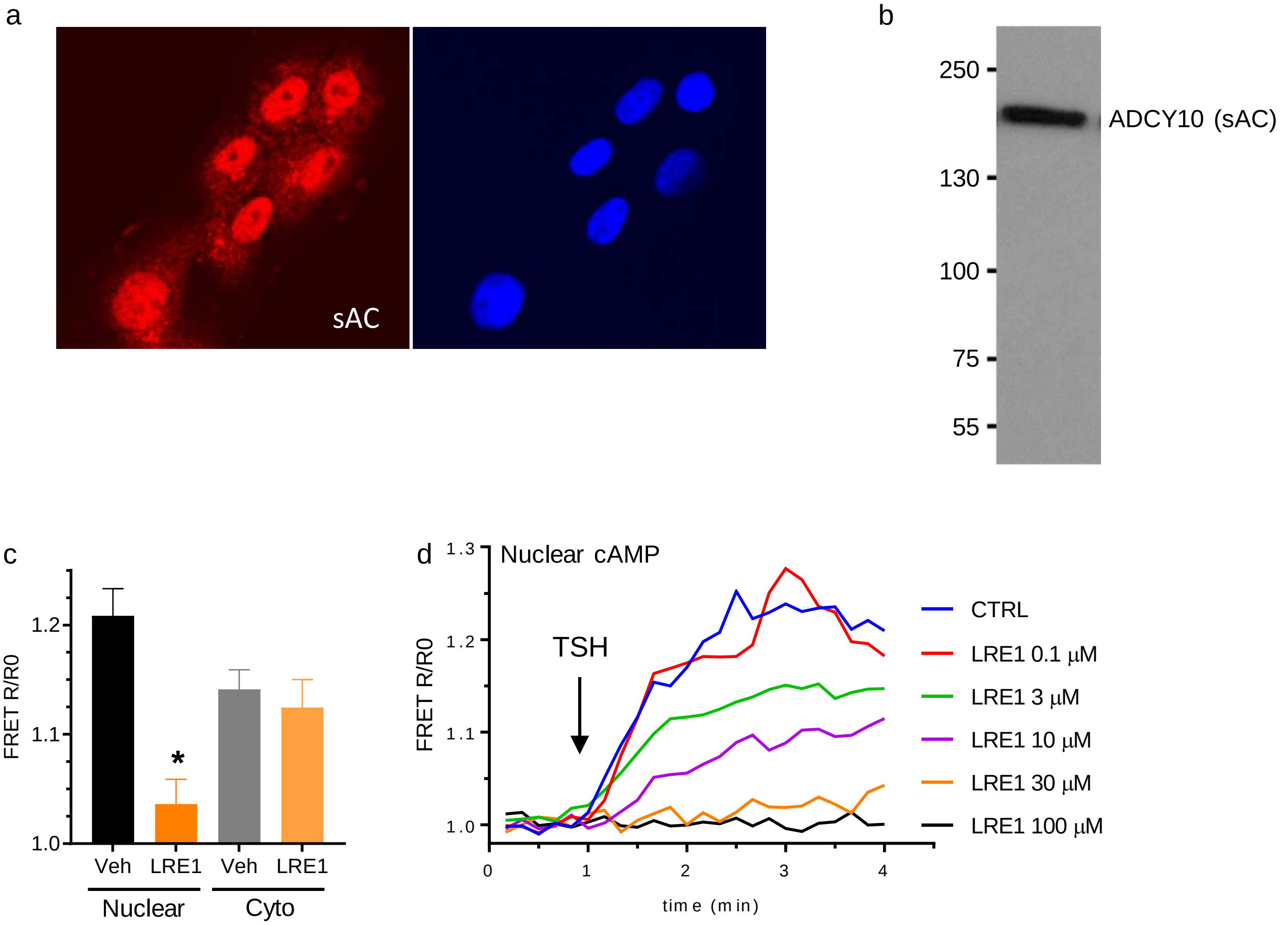


**Fig. S2.** (a) Immunofluorescence staining of endogenous sAC (ADCY10) in PCCL3 cells. Images (XY plane) were acquired using confocal microscopy and sAC is shown in red (left) and DAPI in blue (right). (b) Immunoblotting of endogenous sAC in PCCL3. (c) Normalized ratios (R/R0) of the same experiments show in Fig. 1H. (d) Normalized ratios (R/R0) of representative cell recordings for the experiments shown in Fig. 1I. Data are expressed as mean ± SEM (for n cells/samples and independent experiments, see statistics). Significance was tested using a two-tailed Student's t-test (*, $p<0.05$; n.s., not significant).

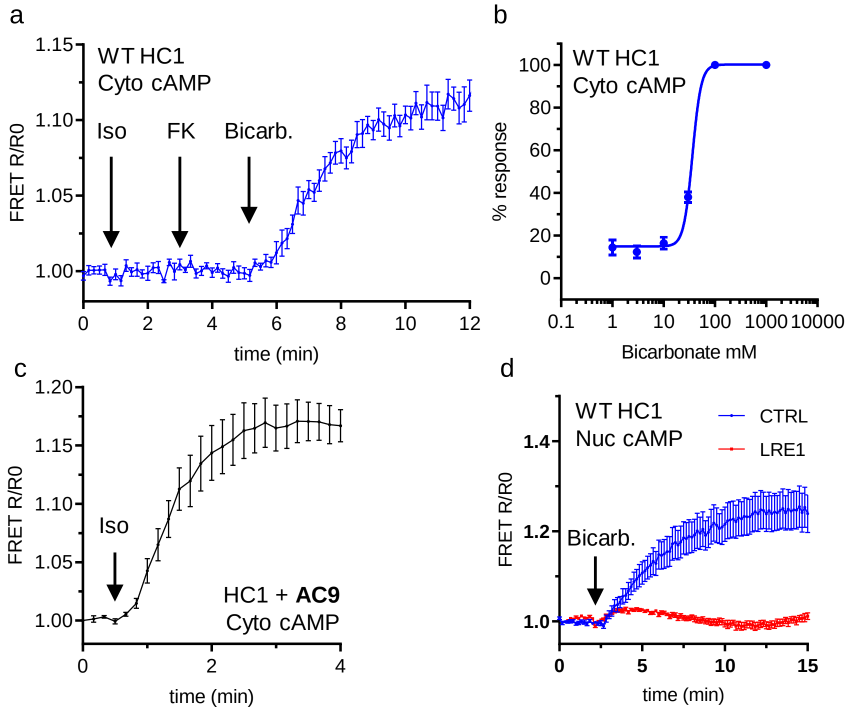


**Fig. S3.** Real-time FRET-based monitoring of cAMP levels in live HC-1 cells. Traces are the normalized FRET ratios (R/R0) of the H208/H188 (cytosolic) or NLS-H208 (nuclear) sensors. (a) Cells transfected with H208 were incubated with Isoproterenol (Iso; 10 μM), Forskolin (25 μM) and Bicarbonate (80 mM). (b) Bicarbonate dose-response experiments performed in HC1 cells transfected with H188. (c) HC1 cells transfected with AC9 and H208 were incubated with Isoproterenol (Iso; 10 μM). (d) HC1 transfected with NLS-H208 preincubated with Veh (CTRL) or LRE1 (100 μM) for 30 min before the introduction of bicarbonate (80 mM). Data are expressed as mean ± SEM (for n cells/samples and independent experiments, see statistics).

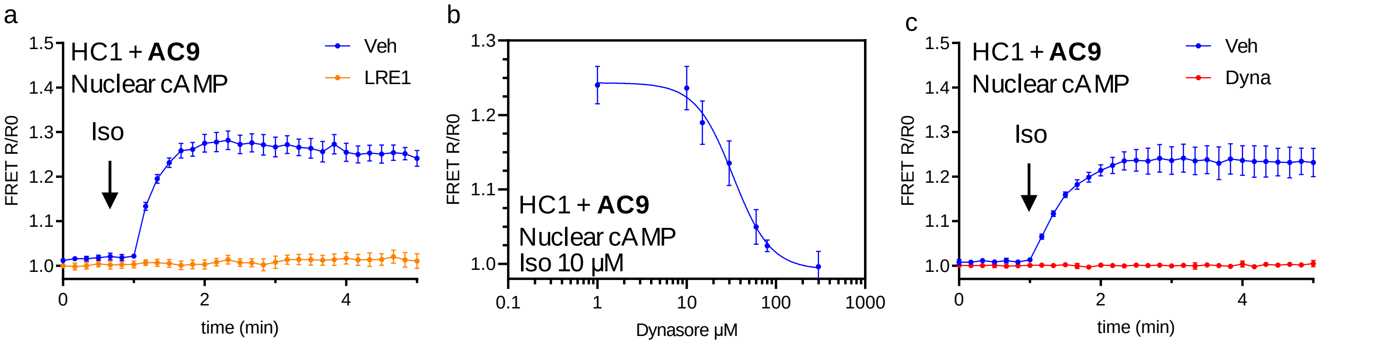


**Fig. S4.** Real-time FRET-based monitoring of cAMP levels in live HC-1 transfected with AC9 cells. Traces are the normalized FRET ratios (R/R0) of the NLS-H208 (nuclear) sensor. (a) Cells were preincubated with Vehicle (Veh) or LRE1 (100 μM) for 30 min before the introduction of isoproterenol (Iso; 10 μM). (b) Concentration-dependent inhibition of the Iso response by Dynasore. (c) Cells were preincubated with Vehicle (Veh) or Dynasore (Dyna; 30 μM) for 15 min before the introduction of isoproterenol (Iso; 10 μM). Data are expressed as mean ± SEM (for n cells/samples and independent experiments, see statistics).

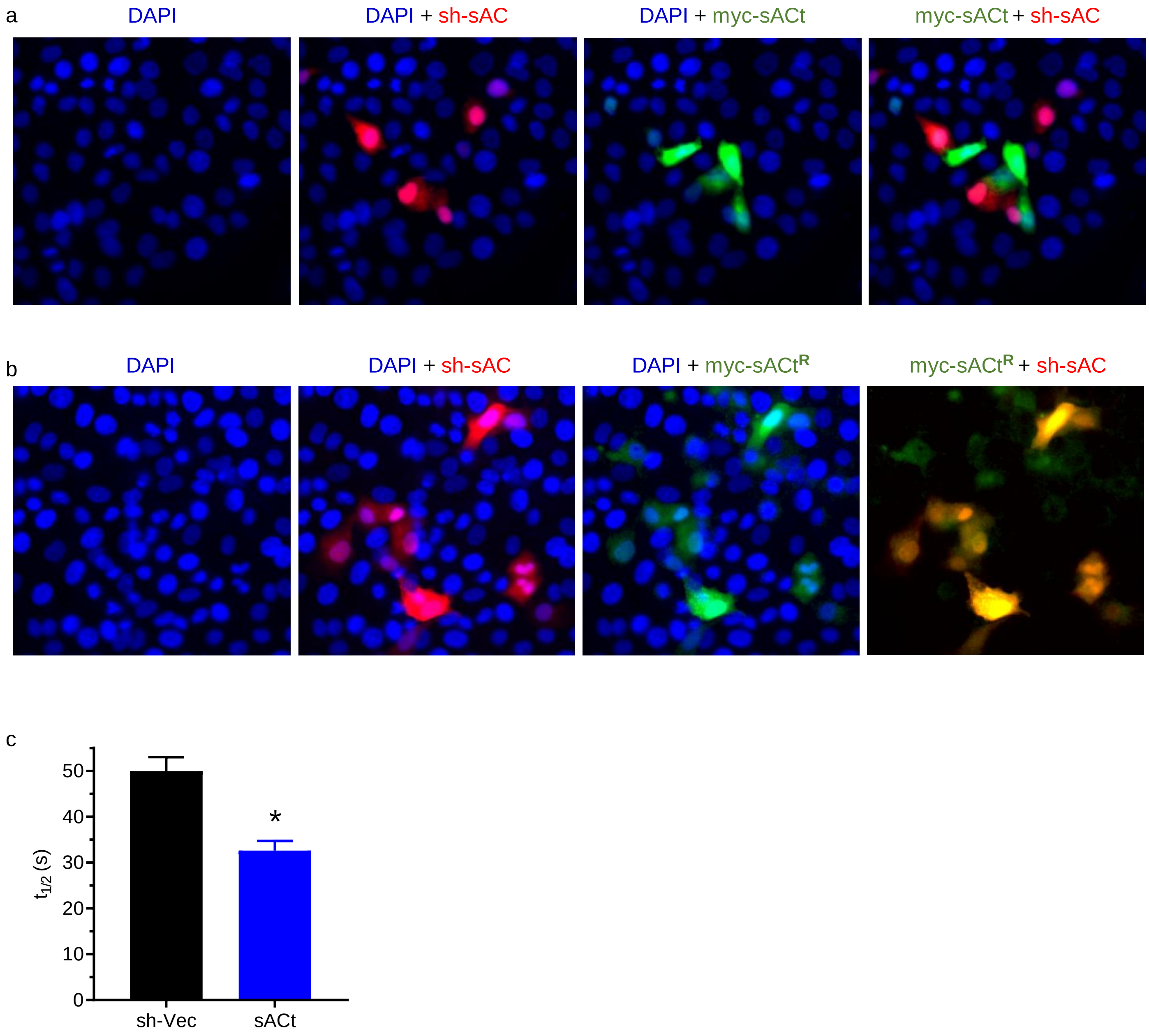


**Fig. S5.** (a) Cells were transfected with pSiren-red-sh-sAC1 (sh-sAC; red) and sACt (myc-sACt; green). (b) Cells were transfected with pSiren-red-sh-sAC1 (sh-sAC; red) and sh-resistant sACt (myc-sACt$^R$; green). Note the absence of co-staining (yellow) with sh-sensitive sAC. (c) Quantification ($t_{1/2}$) of the kinetic traces shown in Fig. 2d. Data are expressed as mean ± SEM (for n cells/samples and independent experiments, see statistics). Significance was tested using a two-tailed Student's t-test (*, $p<0.05$; n.s., not significant).

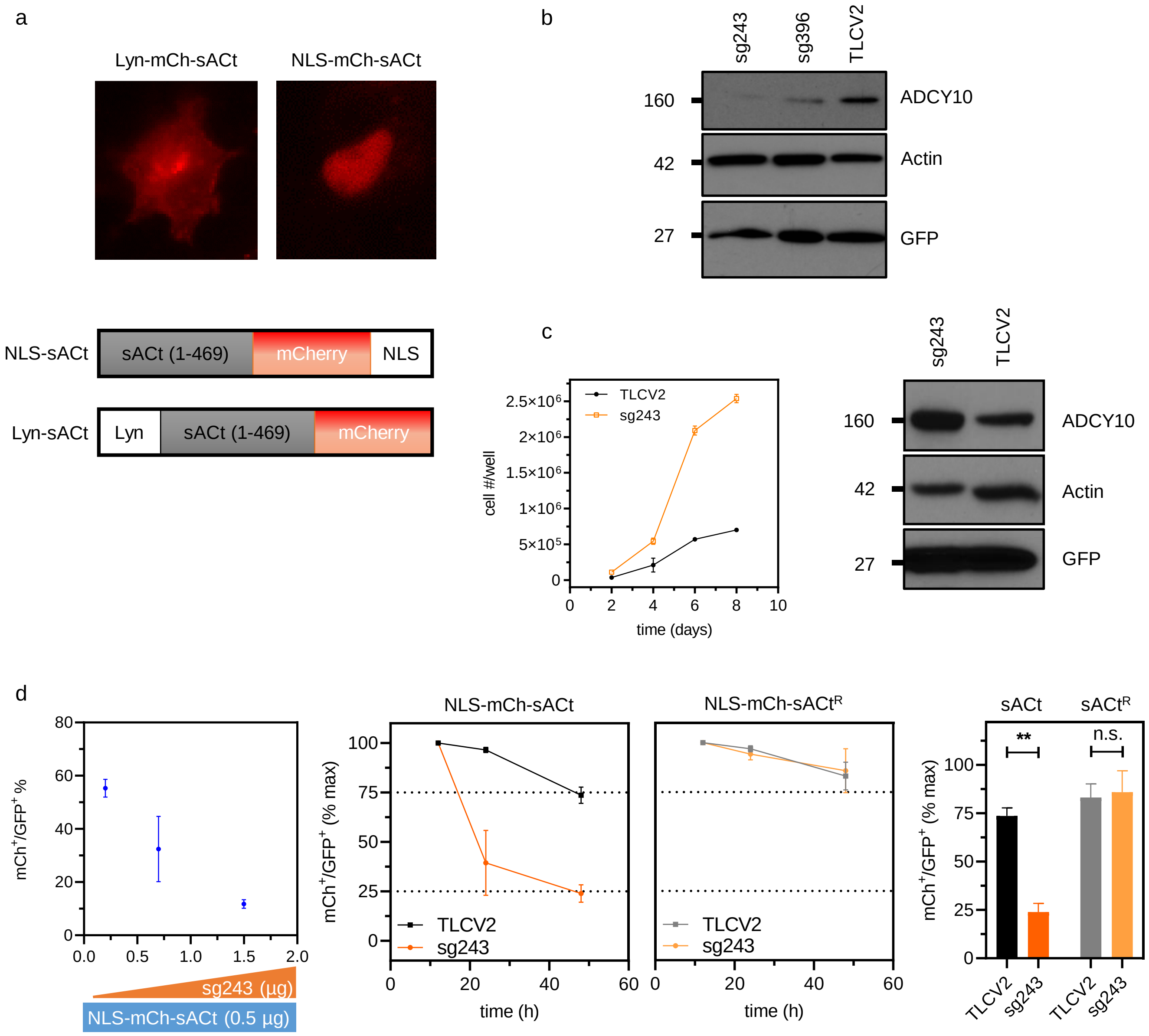


**Fig. S6.** (a) Cellular distribution of Lyn-mCherry-sACt (left) and NLS-mCherry-sACt (right) and their organization (lower cartoon). (b) Transient effect of lentivirus-mediated infection of PCCL3 cell with TLCV2 vector and TLCV2-sg243, or TLCV2-sg396 constructs. Cells were stimulated with doxycycline for three days. The efficiency of sg243 deletion was superior to sg396 and was used in all the studies reported. (c) Upon two weeks of expansion in puromycin, no sAC-KO stable clones were observed (consistent with sAC being required for proliferation) and the population selected shows a higher proliferative rate (left) and with higher sAC expression level (right) (consistent with sAC being rate-limiting for proliferation). (d) Titration experiments showing the effect of increasing amounts of TLCV2-sg243 (doxycycline/GFP$^+$) on the expression levels of a fixed amount of NLS-mCherry-sACt (mCh$^+$; 0.5 µg) after 48h (left panel). Using a 2:1 (TLCV2-sg243:NLS-mCherry-sACt/sACt$^R$) ratio, time-courses confirmed that while NLS-mCh-sACt expression in GFP+ cells decreased over time, sg243-resistant (NLS-mCh-sACt$^R$) levels remained stable (middle panels). Quantification of these data at 48h is shown on the right. Data are expressed as mean ± SEM (for n cells/samples and independent experiments, see statistics). Significance was tested using a two-tailed Student's t-test (**, p<0.01; n.s., not significant).

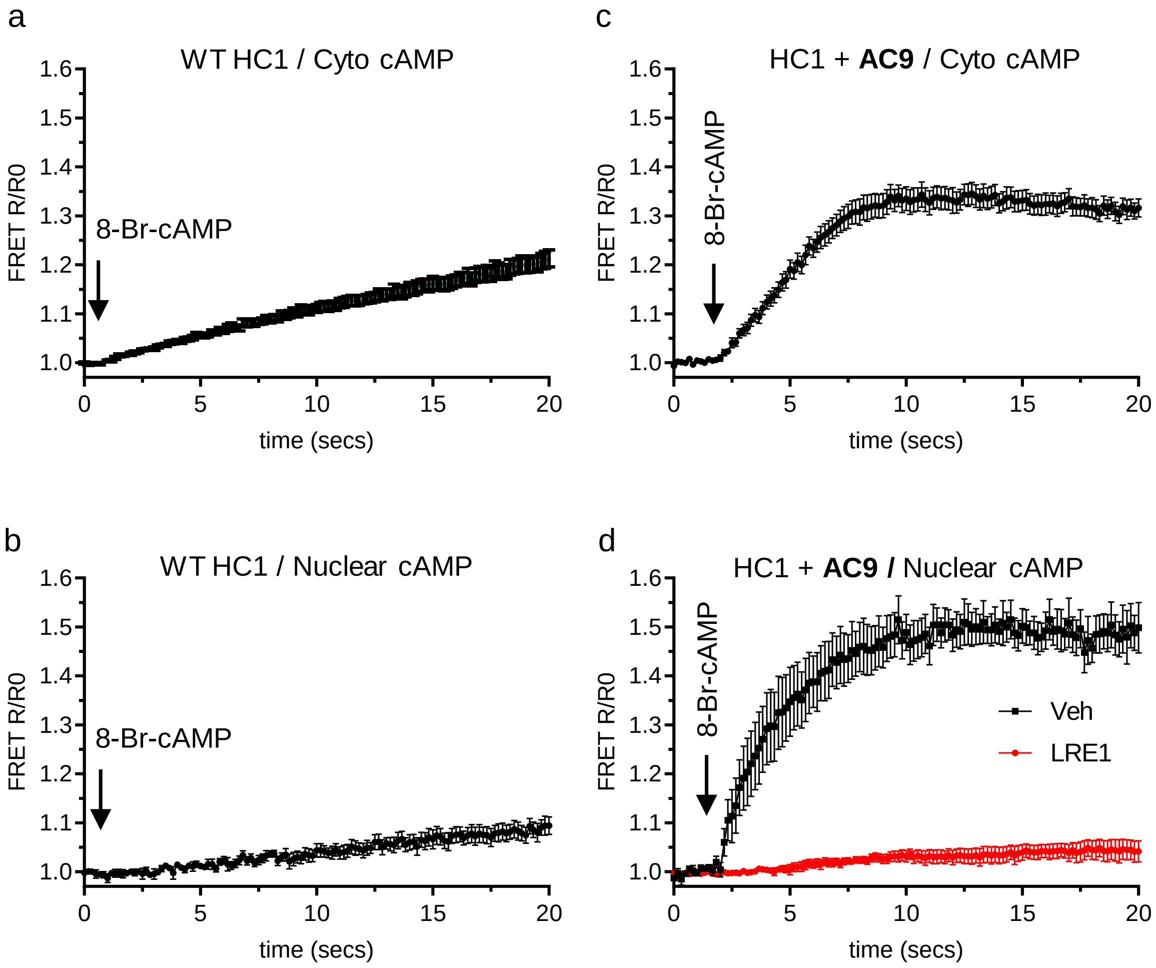


**Fig. S7.** Real-time FRET-based monitoring of cAMP levels in live HC-1 cells and incubated with 8-Br-cAMP (3 mM). Traces are the normalized FRET ratios (R/R0) of cytosolic (H208) or nuclear (NLS-H208) sensors. (a) WT HC1 cells were transfected with H208 (a) or NLS-H208 (b). AC9-transfected cells were cotransfected with H208 (c) or NLS-H208 and preincubated with Vehicle (Veh; DMSO) or LRE1 (100 μM) for 30 min. Data are expressed as mean ± SEM (for n cells/samples and independent experiments, see statistics).

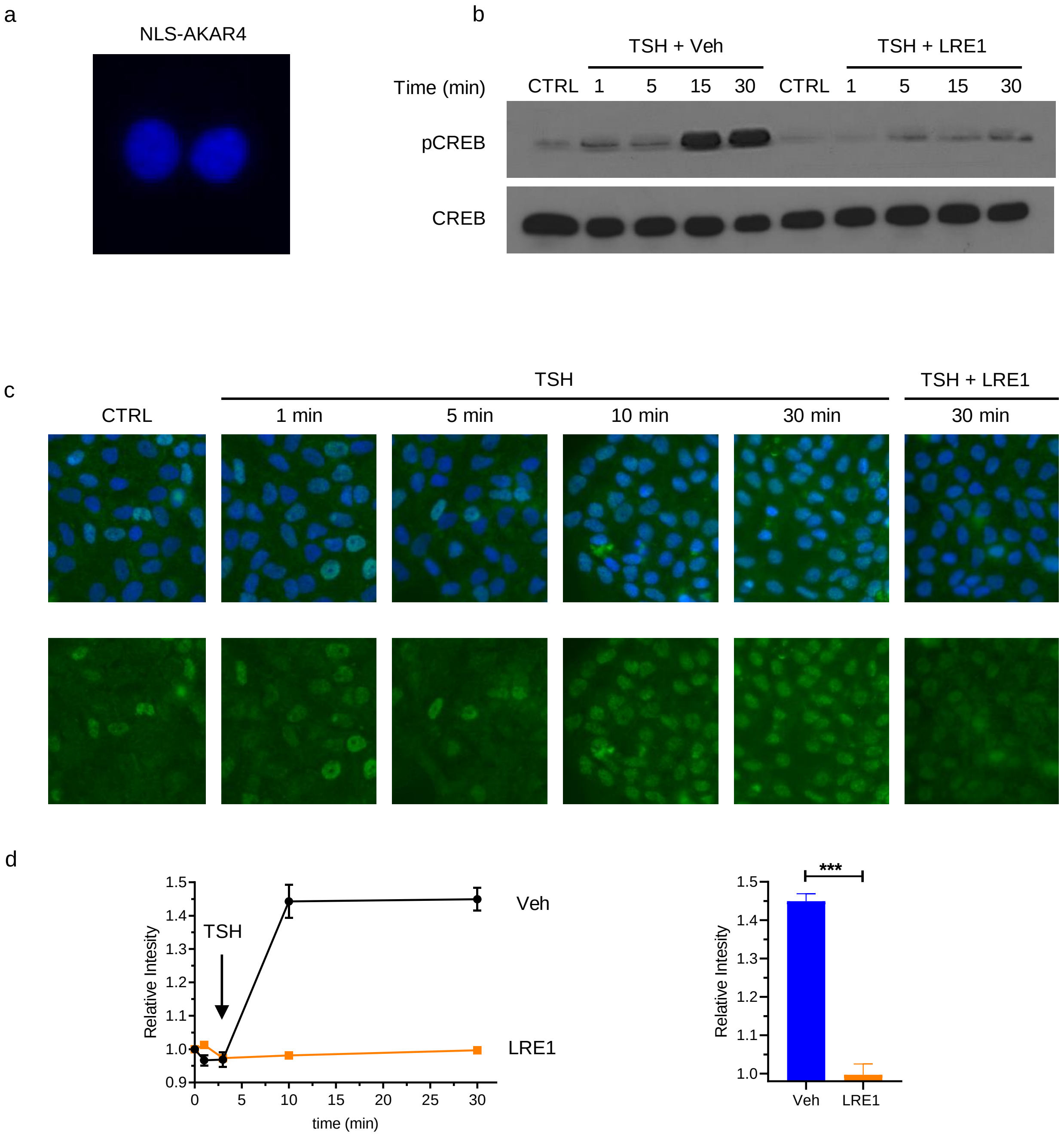


**Fig. S8.** (a) Fluorescence (cyan channel; Cerulean; peak emission, 475nm) confirms the nuclear localization of the NLS-AKAR4 sensor in PCCL3 transfected cells (two nuclei shown). (b) pCREB (upper panel) and total CREB (lower panels) immunoblots of PCCL3 cells in starving conditions for 20h (CTRL; no TSH) or after 1, 5,10, or 30 min of TSH stimulation (1 mIU/ml). Cells were preincubated with vehicle (TSH + Veh) or LRE1 for 30 min (TSH + LRE1). (c) Similar experiments where PCCL3 cells were starved for 20h and stimulated with TSH and fixed and stained with anti-pCREB antibodies and DAPI. (d) Quantification of the experiments in S8c, measuring fluorescence intensity (pCREB channel) normalized to TSH at t=0. Data are expressed as mean ± SEM (for n cells/samples and independent experiments, see statistics). Significance was tested using a two-tailed Student's t-test (***, p<0.001; n.s., not significant).

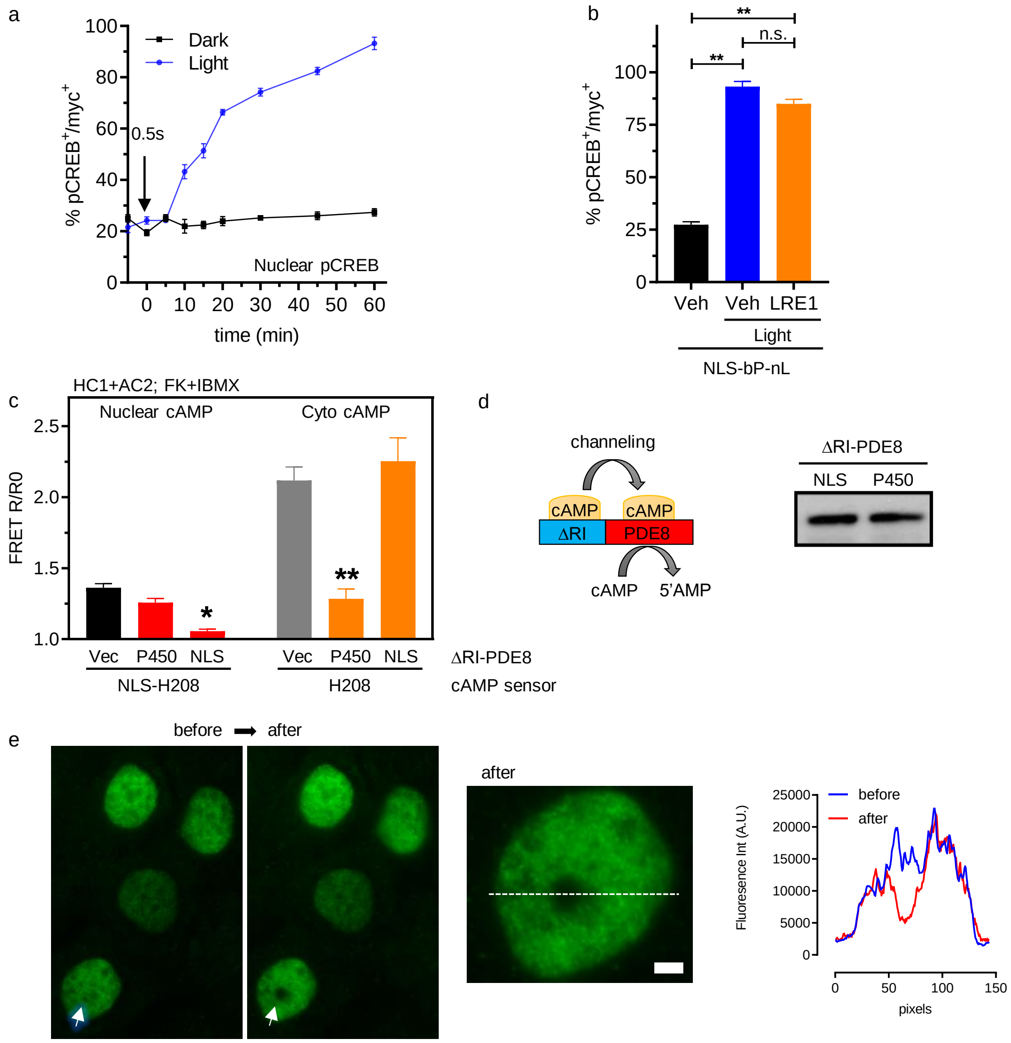


**Fig. S9.** (a) PCCL3 cells stably transfected with NLS-bPac-nLuc (myc+) were stimulated with a single pulse of light (0.5 s) using the custom-built Arduino-controlled LED system (see methods). Cells were fixed at 5,10,15, 20, 30, 45, and 60 min after light stimulation and nuclear CREB phosphorylation was assessed by immunofluorescence. Data is expressed as % pCREB+/myc+ cells. (b) Quantitative analysis of the % of pCREB+/myc+ in non-stimulated and stimulated cells (0.5s, light). Cells were preincubated with Veh or LRE1 (100 μM) for 30 min. (c) HC1 cells were transfected with AC2 and a nuclear or cytosolic FRET-based cAMP sensor (NLS-H208 or H208) and either Vector (Vec), NLS-ΔRI-PDE8, or P450-ΔRI-PDE8 constructs. Cells were stimulated with FK (20 μM) + IBMX (250 μM) to evaluate the effectivity of the targeted ΔRI-PDE8 constructs to eliminate the compartment-specific cAMP response. (d) The cartoon represents the recently described ability of PKA-RIa to enhance the hydrolytic activity of PDE8 via channeling (left). Expression of ΔRI-PDE8 constructs in HC1 cells (FLAG, right). (e) Testing the UGA-42 Geo system for localized photostimulation. Cells were labeled with a nuclear Alexa 488-marker and stimulated with the UGA-42 using a long (30s), laser pulse (445 nm) directed to the nuclear center. The photobleaching area was ~1 μm$^2$ without affecting the surrounding fluorescence. The arrow points to the targeted (nuclear center) area before and after bleaching (left panels). Detail of the bleached nuclei (middle panel; bar = 1 μm). Fluorescence intensity (A.U.; arbitrary units) along the line scan shown in the middle panel (right panel). Data are expressed as mean ± SEM (for n cells/samples and independent experiments, see statistics). Significance was tested using a two-tailed Student's t-test (*, p<0.05; **, p<0.01; ***, p<0.001; n.s., not significant).

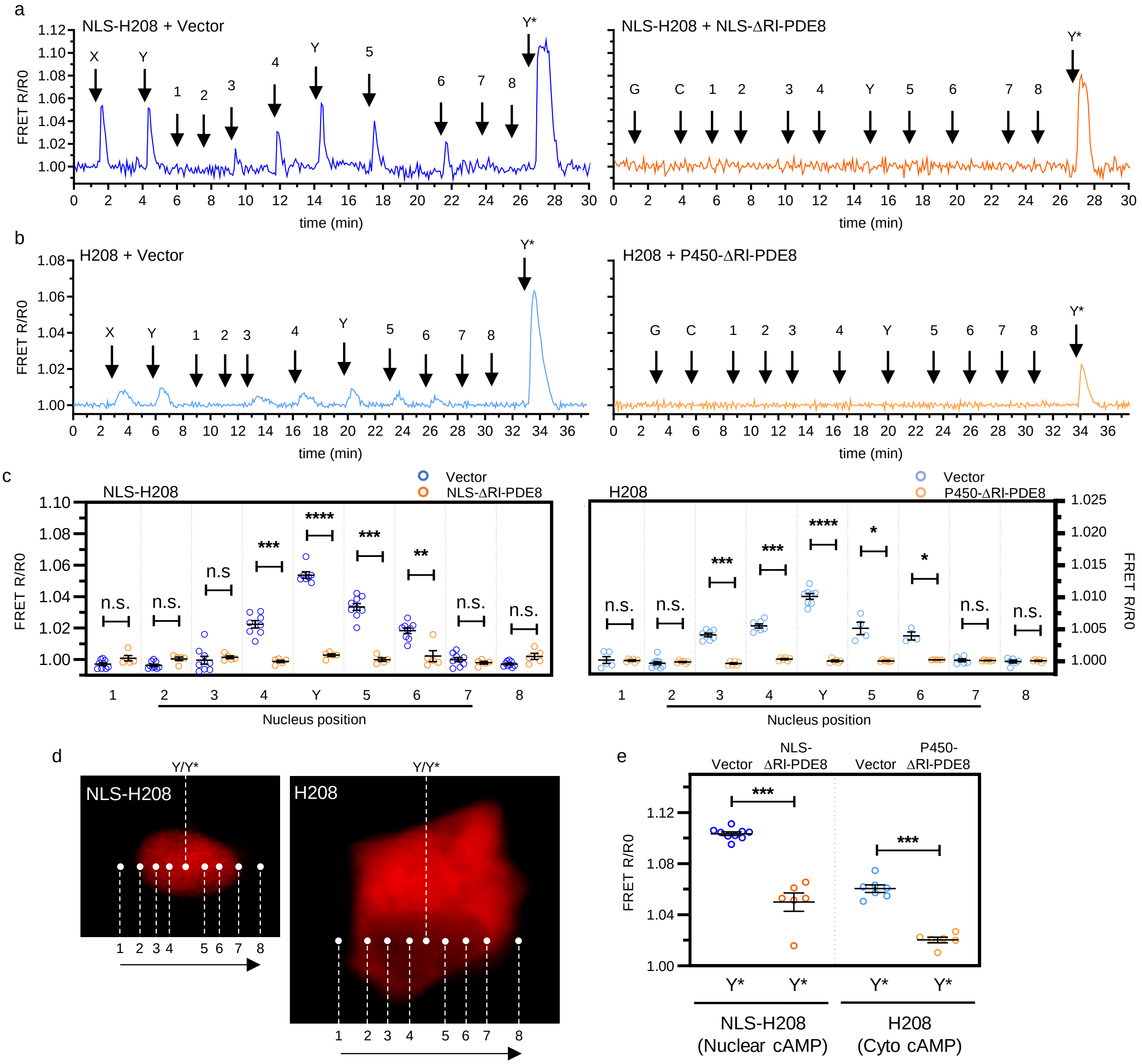


**Fig. S10.** PCCL3 cells stably transfected with NLS-bPac-nLuc (NLS-bP-bL) were transiently transfected with the nuclear (NLS-H208) or cytosolic (H208) cAMP sensors. Cells were photostimulated with a 445 nm laser controlled by the UGA-42 point-scanning system that allowed us to stimulate predetermined small areas (~1 μm). Stimulation parameters (pulse train: 50 ms, 300 ms total duration, 10 Hz, 3% laser power, 1% physical ND filter) were set empirically to match the cAMP signal generated by the Arduino-controlled LED system that was used in the proliferation experiments (0.5s, 20% LED power; see Fig. 6). While the LED system produces global stimulation (X), generating a cAMP signal in all cells expressing NLS-bP-bL, the point scanner system allowed us to target and stimulate a small part of the nuclear center (Y) in a single cell using a small ROI. After checking the similarity of the X and Y responses, cells were sequentially stimulated with the same potency along a straight line (1, 2, 3, 4, Y, 5, 6, 7, 8; see panel d) passing through the nuclear center, visualized by the sensor fluorescence. Stimulations 1 and 8 were targeted outside the nuclei, 2 and 7 were targeted at the nuclear edge,

3 and 6 were targeted at an intermediate position and 4 and 5 were targeted right next to the nuclear center (Y). At the end of each experiment, a higher power stimulation (single 30s pulse, 30% laser power, 1% physical ND filter) targeted to the nuclear center (Y*) was performed to rule out sensor saturation in any of the previous stimulations. Representative traces (normalized ratios; [FRET R/R0]) of individual cells transfected with NLS-H208 and Vector (a, left panel) or NLS-ΔRI-PDE8 (a, right panel), and in cells transfected with H208 and Vector (b, left panel) or NLS-ΔRI-PDE8 (b, right panel). These experiments showed that ΔRI-PDE8 constructs were able to inhibit the cAMP elevation generated by a low-intensity stimulation (Y), equivalent to that used in proliferation studies. In addition, these experiments showed that cAMP is generated only when stimulation is directed to the nucleus indicating that there is no off-target expression of NLS-bPAC-nLuc. Note that pulses targeted to the nuclear edge were unsuccessful to produce a detectable cAMP signal, suggesting that a minimum activity of NLS-bPAC-nLuc has to be stimulated to produce a detectable signal. Although a high-intensity stimulation (Y*) generated a cAMP elevation in both compartments, even in the presence of the ΔRI-PDE8 constructs, the cAMP response was significantly diminished. (c) Quantitative analysis of the stimulation site vs. cAMP response (FRET R/R0) in the nucleus (NLS-H208; left panel) or cytosol (H208; right panel) in cells transfected with Vector or one of the ΔRI-PDE8 constructs. (d) White dots indicate the places where cells were stimulated; from left to right: 1, 2, 3, 4, Y, 5, 6, 7, 8. While “Y” denotes a low-power stimulation, “Y*” denotes a high-power stimulation that was directed to the nuclear center. (e) Quantitative analysis of the stimulation site vs. nuclear (NLS-H208) or cytosolic (H208) cAMP response (FRET R/R0) of cells due to a high-power stimulation directed to the nuclear center (Y*) in cells transfected with Vector or one of the ΔRI-PDE8 constructs. Data are expressed as mean ± SEM (for n cells/samples and independent experiments, see statistics). Significance was tested using a two-tailed Student's t-test (*, $p<0.05$; **, $p<0.01$; ***, $p<0.001$; n.s., not significant).
